# An integrated genome-wide resource reveals distinct replication environments of DNA breakage in human cancer cell lines

**DOI:** 10.64898/2026.09.10.750571

**Authors:** Boyu Ding, Alex Ing, Li-Chin Wang, Lorenzo Corazzi, Marco Giaisi, Victoria Marini, Lumír Krejčí, Diala Shatleh, Rami Aqeilan, Pei-Chi Wei

## Abstract

Replication stress is a major source of genome instability in cancer, yet the genomic features that determine where DNA double-strand breaks (DSBs) arise remain incompletely defined. Here, we establish an integrated genome-wide resource of DSBs, DNA replication, replication timing, transcription, and R-loops in two widely used cancer cell lines, U2-OS and HeLa, under steady-state conditions and following prolonged low-dose DNA polymerase inhibition. We combine these datasets with systematic statistical testing and comparative analytical approaches to define the replication environments associated with genome fragility. Endogenous DSBs preferentially accumulated at origin-rich initiation zones, where R-loops were enriched, whereas prolonged DNA polymerase inhibition redirected DSB formation toward origin-poor, late-replicating regions. Among the replication features examined, replication initiation zones and late-replicating areas were most sensitive to prolonged replication stress. R-loops were specifically enriched at initiation zones but depleted from late-replicating regions and recurrent DNA break clusters (RDCs), demonstrating that their association with genome fragility is context dependent. Replication stress further induced RDCs within long, actively transcribed genes, while their locations only partially overlapped with common fragile sites. Functional analyses identified MUS81 as a major regulator of RDC formation. MUS81 loss increased RDCs, whereas restoration of its catalytic activity suppressed them, indicating that MUS81 resolves replication intermediates before they persist into late-replicating fragile regions. Together, this resource and analytical framework provide a systematic basis for dissecting how replication architecture, transcription, and DNA processing shape genome fragility in cancer cells.

## Introduction

Genome instability is a hallmark of cancer, and a major driver of tumor evolution(1). Because cancer cells divide rapidly, they put increased demands on DNA replication and repair machinery, which can increase the likelihood of errors. These errors can lead to genomic rearrangements, such as translocations, which bring enhancers or promoters into proximity with oncogenes, thereby acting as powerful drivers of tumorigenesis. A major source of these errors is replication stress, where stalled replication forks undermine genome stability. Transcription–replication conflicts (TRCs) are major sources of intrinsic replication stress. They occur when replication forks collide with DNA replication and transcription machinery, or from the formation of DNA:RNA hybrids. When TRCs occur, replication forks stall and undergo remodeling. Stalled forks can be either cleaved by nucleases such as MUS81, or undergo fork reversal, forming newly synthesized, single-ended DNA ends. Both fork cleavage and fork reversal therefore yield single-ended DNA substrates prone to chromosomal translocation (2). As long-range rearrangements can juxtapose oncogenes with enhancers (3) or disrupt tumor suppressor genes (4). Defining the origin of these unstable DNA ends is essential for understanding how replication stress promotes genome instability in cancer.

We previously showed that ectopic replication stress in non-cancerous neural progenitor cells gives rise to recurrent DNA break clusters (RDCs) (5–7). RDCs are genomic regions where DNA double-strand breaks (DSBs) occur recurrently at defined positions across cells, identified directly by genome-wide DSB mapping. RDC occurs preferentially in actively transcribed genes - over 70% of which exceed 300 kb - features that render them particularly vulnerable to TRCs. A subset of RDCs occurred at early-replicating loci enriched for DNA:RNA hybrids, whereas ∼30% localized to late-replicating regions overlapping with common fragile site (CFS) (6), which are classically defined by cytogenetic fragility and under-replication upon replication stress. Despite this substantial overlap, RDCs and CFSs are not equivalent: RDCs provide a direct molecular map of recurrent break formation, whereas CFSs represent genomic regions prone to replication failure and manifest as chromosomal gaps or breaks during mitosis (5). Thus, RDCs reflect DNA breaks arising at stalled replication forks, indicating that their origin precedes CFS expression and occurs during S phase. Importantly, RDCs outside of early-replicating regions consisted of single-ended DNA breaks aligned with the direction of replication, implicating its potential origins as nuclease-mediated cleavage of stalled forks. Together, these findings suggest that distinct classes of RDCs likely reflect cell type– specific replication and transcription programs, underscoring the need for comprehensive, matched datasets to define their origins across different cellular contexts.

Indirect evidence suggests a role for RDCs in cancer mutagenesis. A subset of RDCs coincide with tumor suppressor genes, suggesting that their disruption could promote malignant transformation (5, 6). Others colocalize with structural variation hotspots at long-noncoding RNA loci (e.g. PVT1, NEAT1) that drive tumor progression (8, 9), indicating that RDCs may serve as initiation sites for oncogenic rearrangements. Furthermore, a cancer-associated copy number signature, CN9, exhibits features consistent with RDC-mediated DNA breaks (10), reinforcing their potential relevance to cancer genesis. Nevertheless, RDCs have, so far, only been characterized in neuronal lineages (5, 6, 11) and murine embryonic stem cells (12). Although similar structures occur in other cell types, cancer cells are inherently burdened by replication stress. Yet, resistance often emerges as cancer cells adapt their stress responses and shift damage-prone sites, underscoring the need to understand how replication stress reshapes the DNA damage landscape.

To address these questions, we generated matched genome-wide maps of DNA replication, replication timing, transcription, DNA hybrids, and DSBs in U2-OS, HeLa, and HeLa-Kyoto cells under steady-state conditions and following prolonged DNA polymerase inhibition. Integrative analyses revealed distinct replication environments of endogenous and stress-induced DSBs – endogenous breaks preferentially localized to replication initiation zones enriched for R-loops , whereas polymerase inhibition redirected breaks to origin-poor, late-replicating regions. We further identified RDCs within long, actively transcribed genes, which were depleted of R-loops, indicating distinct determinants of genome fragility across replication environments. Functional analyses identified MUS81 as a regulator of RDC formation. Together, these datasets and analytical approaches provide a resource for defining how replication architecture, transcription, and DNA processing shape genome fragility in human cell lines.

## Materials and Methods

### Cell culture

U2-OS and HeLa cells were provided by Dr. Chong Sun (Division of Immune Regulation in Cancer, German Cancer Research Center, Heidelberg, Germany) and Dr. Haikun Liu (Division of Molecular Neurogenetics, German Cancer Research Center, Heidelberg, Germany), respectively. A panel comprising parental U2-OS WT, MUS81-KO and GEN1-KO cells was obtained from the laboratory of Prof. Stephen C West (DNA Recombination and Repair Laboratory, The Francis Crick Institute, London, UK). These knockout cell lines were generated and validated as previously described (13). Parental HeLa-Kyoto WT and MUS81-KO cells were provided by Prof. Pavel Janscak (Institute of Molecular Cancer Research, University of Zurich, Zurich, Switzerland) and were previously described by Chappidi et al. (14). HeLa-Kyoto represents a distinct HeLa-derived cell-line background from the HeLa cells used for the genome-wide comparative analyses (15). The identities of all cell lines were authenticated using the Multiplex human Cell line Authentication Test (Multiplexion GmbH, Heidelberg, Germany). Cells were maintained in Dulbecco’s modified Eagle’s medium supplemented with 10% fetal bovine serum, penicillin, streptomycin and L-glutamine at 37°C in a humidified atmosphere containing 5% CO₂. Detailed reagent information is provided in Supplementary Methods.

### MUS81 reconstitution and siRNA-mediated depletion

For transient MUS81 reconstitution, U2-OS MUS81-KO cells were co-transfected with the chromosome 1 LAM-HTGTS bait-inducing plasmid and either pAIO-MUS81^WT^ or pAIO-MUS81^DD/AA^, encoding N-terminally EGFP-tagged MUS81 proteins, at a bait-to-expression-plasmid mass ratio of 3:1. Transfection was performed using polyethyleneimine (PEI) at a total DNA-to-PEI mass ratio of 1:1. Following transfection, cells were treated with 0.4 μM aphidicolin or left untreated for 72 h. Three days after transfection, EGFP-medium and EGFP-low/negative populations were isolated by fluorescence-activated cell sorting. EGFP-medium cells were used for LAM-HTGTS, cell cycle analysis and Western blot, whereas parallel EGFP-low/negative populations were collected for Western blot.

For MUS81 knockdown, U2-OS cells were transfected with 20 nM siRNA -MUS81 or Silencer Select Negative Control No. 1 siRNA. Cells were subsequently transfected with the chromosome 1 LAM-HTGTS bait plasmid and treated with 0.4 μM aphidicolin or left untreated for 72 h. Parallel samples were collected for Western blot to assess MUS81 protein level. Detailed transfection and validation procedures are provided in Supplementary Methods.

### Construction of the pAIO-MUS81^DD/AA^ plasmid

The pAIO-MUS81^DD/AA^ plasmid, encoding the MUS81 D338A/D339A mutant, was generated from pAIO-MUS81^WT^ (16) by site-directed mutagenesis. The complete plasmid sequence was examined by Oxford Nanopore sequencing, and the intended D338A/D339A substitutions were independently confirmed by Sanger sequencing. Primer sequences and detailed cloning procedures are provided in Supplementary Methods.

### High resolution Repli-seq (HiRepli-Seq)

High-resolution 16-fraction Repli-seq was performed essentially as previously described (6,13), with minor modifications. Cells were treated with 0.4 μM aphidicolin or left untreated for 16 h. Nascent DNA was labelled with 400 μM BrdU for 30 min in untreated cells or 45 min in aphidicolin-treated cells. Ethanol-fixed cells were stained with propidium iodide, and S-phase cells were separated by flow cytometry into 16 consecutive fractions, with at least 60,000 cells collected per fraction.

Genomic DNA was extracted from the sorted cells, fragmented to an average size of approximately 200 bp, converted into sequencing libraries and subjected to anti-BrdU immunoprecipitation. Libraries were sequenced on an Illumina NextSeq using 75-bp single-end chemistry or on an Illumina NovaSeq X Plus using 100-bp paired-end chemistry. Matched G1-phase genomic DNA was collected and subjected to whole-genome sequencing for DNA-copy-number normalization. Detailed HiRepli-seq library preparation, G1 sorting, data processing, scaling and replication-feature calling procedures are provided in Supplementary Methods.

### Linear amplification-mediated, High-throughput, genome-wide translocation sequencing (LAM-HTGTS)

To induce targeted bait sites on chromosomes 1, 3, 11, approximately 4 × 10⁶ cells were transfected with 20 µg of the spCas9/sgRNA expression plasmid (pX330-U6-Chimeric-BB-CBh-hSpCas9) using PEI at a 1:1 DNA:PEI ratio (20 µg DNA/ 20 µg PEI). SgRNA sequences specific to each bait site were individually cloned into pX330 vectors following the protocol described by the Zhang lab (https://www.addgene.org/crispr/zhang/). and their genomic coordinates and oligonucleotide sequences are provided in Supplementary methods.

Following transfection, the cells were subsequently treated with 0.4 µM aphidicolin or left untreated for 96 hours. Following treatment, cells were harvested, and genomic DNA was extracted with a standard phenol/chloroform/isopropyl protocol. 10 µg genomic DNA were used as input for library construction and the construction of HTGTS libraries were described before (5, 6, 17). The libraries were sequenced using an Illumina NextSeq 550. For the MUS81 reconstitution and siRNA-depletion experiments performed using the chromosome 1 bait, cells were treated with DMSO or 0.4 μM aphidicolin for 72 h. Detailed bait information and library-construction procedures are provided in Supplementary Methods.

Raw reads were demultiplexed with the HTGTS Docker workflow (https://github.com/brainbreaks/HTGTS). Adapters were then removed from the demultiplexed FASTQ files. The cleaned reads were aligned to the human reference genome (hg38/GRCh38) using Bowtie2. Post-alignment processing and library aggregation were performed with the pipeline’s TranslocWrapper, and the resulting result.tlx files containing bait and prey junction information for each library were used for downstream analyses. In untreated cells we detected ∼50,000 DSBs in U2-OS and ∼24,000 in HeLa; aphidicolin treatment raised these numbers to >74,000 and 50,000, respectively. For MUS81 and GEN1 knockout experiments, we recovered ∼29,000 DSBs from aphidicolin-treated parental and MUS81^KO^ U2-OS cells, and ∼20,000 DSBs from the HeLa-Kyoto sets.

Library secondary processing described here was performed with R scripts at the HTGTS-TLX-processing GitHub repository (https://github.com/brainbreaks/HTGTS-TLX-processing). To avoid artefactual signals arising from problematic genomic regions, for all result.tlx files, rows containing junctions fall within the hg38 ENCODE blacklist were removed using the 1.clean_tlx_blacklist.R script. These regions are known to produce spurious high signals due to repetitive sequences, assembly errors, or unannotated repeats, and their removal improves the accuracy of functional genomic assays (18). The cleaned result files were used for downstream analyses. For statistical analyses and visualization (Figures 3, 4), we exclude the breaks on the bait chromosome (2.TLX_statistics.R function.R) as the bait preferentially recovers many more DSBs at the break site chromosome than the other non-bait chromosome. For visualizing DSB density across the hg38 genomes, we applied functions in the 3.extend_normalize_split.R script, creating BED files with normalized junction across experimental conditions, in which the junction was extended 50kb at each side, producing a 100 kb seed per junction. These seeds were subjected to bedtools genomecov function, converted to bedgraph for IGV graphic views. RDC calling, Analysis of RDC density across genotypes, distance between DSBs to feature-of-interests, microhomology and bait-length usage were described in the Supplementary Materials.

### Global run-on sequencing (GRO-seq)

Global run-on sequencing (GRO-seq) was performed essentially as previously described (5, 6, 12), with minor modifications. Nuclei were isolated from U2-OS or HeLa cells and subjected to nuclear run-on for 5 min at 30°C in the presence of BrUTP. BrUTP-labelled nascent RNA was affinity-purified, converted into strand-specific sequencing libraries and sequenced on an Illumina NextSeq 550 using 75-bp single-end chemistry. Two replicate libraries were generated for each applicable condition. Detailed nuclei-isolation, buffer and library-construction procedures are provided in Supplementary Methods.

GRO-seq libraries were sequenced under NextSeq 550 with a 75 bp single-end high output chemistry. FASTQ files were aligned to the genome build hg38 through Bowtie2 with the docker container GRO-seq pipeline (https://github.com/brainbreaks/GROseq). Strand-specific normalized alignment signal density was converted to bigwig files and reads per thousand per megabase (RPKM) were calculated per refGene entry as described before(6).

### DRIP-seq

DNA:RNA immunoprecipitation sequencing (DRIP-seq) was performed in DMSO-treated U2-OS cells and MUS81-KO cells essentially as previously described (19), with minor modifications. Genomic DNA was extracted and digested with HindIII, EcoRI, XhoI, BsrGI and SspI. DNA:RNA hybrids were immunoprecipitated using the S9.6 antibody. Parallel DNA aliquots treated with RNase H before immunoprecipitation served as specificity controls. Two independent experiments were performed for RNase H-treated and untreated samples. Enrichment at five previously described genomic loci was examined by quantitative PCR before library preparation.

DRIP-seq libraries were prepared using the NEBNext Ultra II DNA Library Prep Kit and sequenced on an Illumina NextSeq 550 using 75-bp single-end chemistry. Reads were aligned to hg38, and significant DRIP-seq enrichment was identified using MACS2. Detailed DNA digestion, immunoprecipitation, library preparation and quality-control procedures were provided in Supplementary Methods.

### Statistical Testing

Details on the various statistical analyses conducted in this investigation are given below. Raw *p*-values for each test are either mentioned in the text, legend (Figures 2B,D, Figures 4C,E, and Figure 5D) , or listed in Table S1. The number of replicates was as follows: GRO-seq (two replicates per condition), LAM-HTGTS (three to eight biological replicates per condition, dependent on the experiments), HiRepli-seq (two replicates per condition), and DRIP-seq (two replicates per condition).

**Figure 1.**
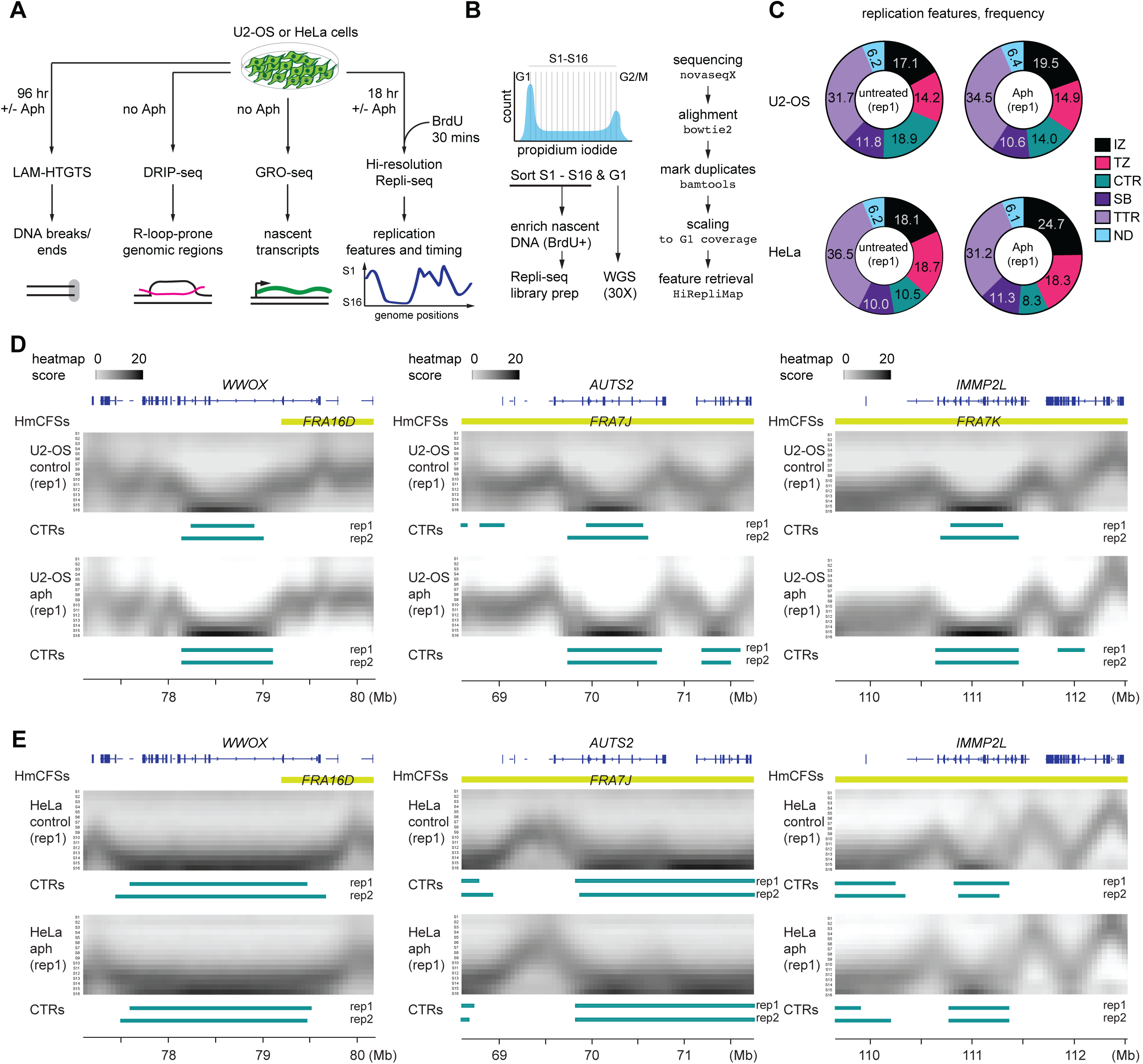
Replication features in U2-OS and HeLa cells. **(A)** Study design and experimental workflow. U2-OS or HeLa cells were subjected to LAM-HTGTS or high-resolution Repli-Seq either with or without aphidicolin (Aph) treatment. For GRO-seq and DRIP-seq, cells were analyzed directly without being treated with Aph. BrdU labeling was used to enrich nascent DNA (HiRepli-Seq) or RNA (GRO-seq), which was then enriched and subjected to protocol-specific library preparation steps. For high-resolution Repli-seq, replication features were identified with HiRepliMap. **(B)** Schematic of experimental design. Cells were BrdU-labeled, sorted into 16 S-phase fractions (S1–S16) plus G1, and subjected to Repli-Seq library preparation and whole-genome sequencing as described in the Materials and Methods. **(C)** Frequency of replication features in untreated versus aphidicolin (Aph)-treated U2-OS and HeLa cells. Pie charts indicate the proportion of initiation zones (IZ), termination zones (TZ), constant timing regions (CTR), steady breakage regions (SB), and timing transition regions (TTR). Percent per feature is indicated. Permutation based p-values quantifying timing transitions between conditions are displayed in Table S1 **(D)** Genome browser views of selected CFSs in U2-OS cells (*WWOX/FRA16D*, *AUTS2/FRA7J*, and *IMMP2L/FRA7K*). Replication timing profiles are displayed as gray heatmaps with late constant timing region (CTR) annotations shown for technical replicates under control and Aph conditions. HmCFS: known fragile sites mapped to human cells’ genomes. **(E)** Genome browser views of the same loci in HeLa cells. Replication timing and CTR annotations are shown as in (D). CTRs at CFS-associated loci remain stable despite global loss of CTRs under Aph treatment.

**Figure 2.**
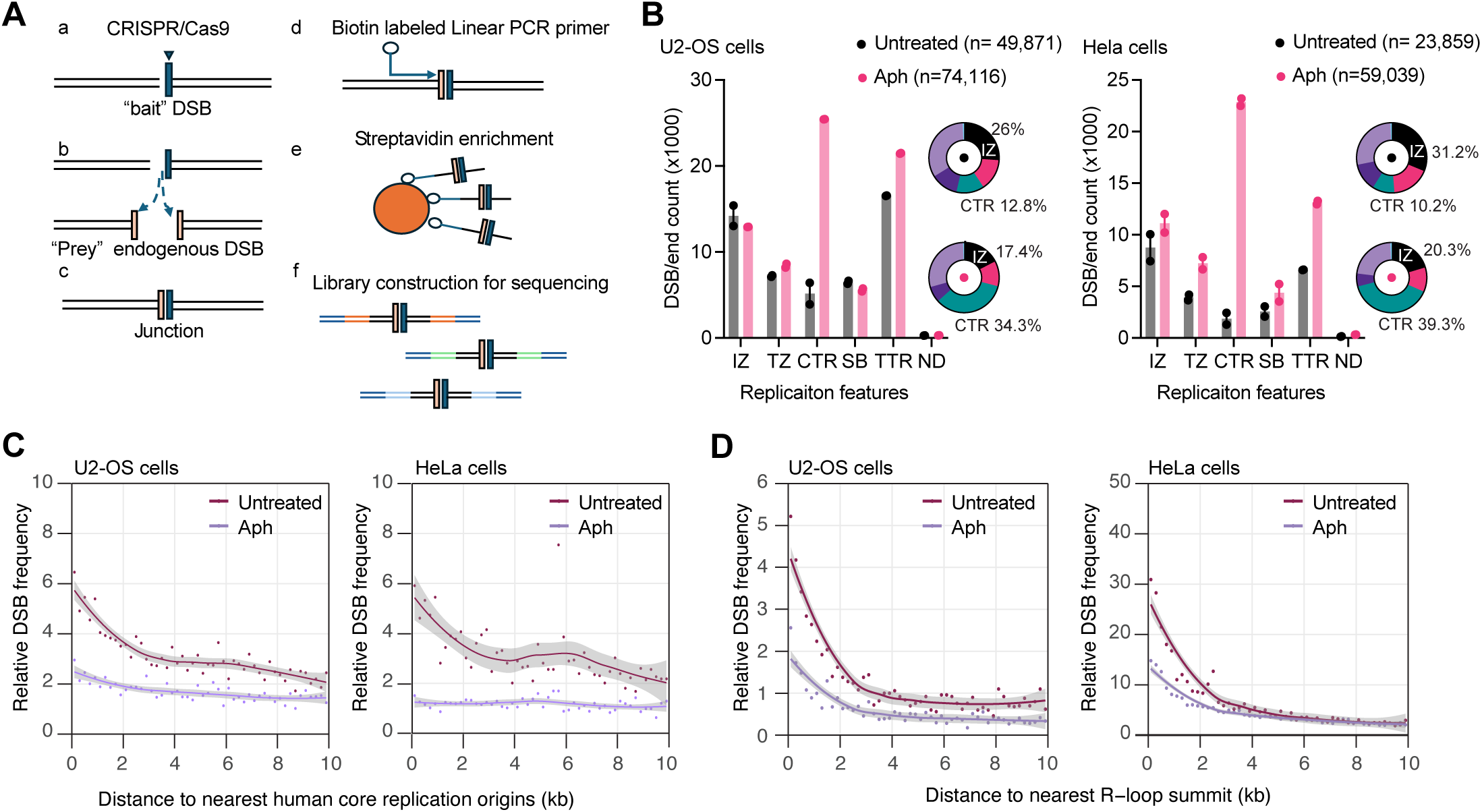
Distribution of DSBs across DNA replication features and correlation with replication origins and R-loops. **(A)** Schematic of the LAM-HTGTS workflow. A CRISPR/Cas9 “bait” DSB captures endogenous “prey” DSBs, which are enriched by linear-amplification–mediated PCR and enriched for library preparation, as described in Materials and Methods. **(B)** Bar plots showing DSB counts and proportions of captured DSB ends mapped to replication features (IZ, TTR, CTR, TZ, SB) in U2-OS and HeLa cells treated with aphidicolin (pink) or left untreated (black). In untreated cells, more than a quarter of all DSB ends localized to IZs, significantly exceeding the genomic coverage of these regions (p < 0.0005, permutation test); no other replication feature was significantly enriched. Upon aphidicolin treatment, breakage shifted predominantly to late-replicating CTRs (p < 0.0005, permutation test). **(C)** Line plots with scatter points showing relative DSB frequency as a function of distance to the nearest human core replication origin (26). lines and shading indicate LOESS fit ± 95% confidence interval. **(D)** Line plots with scatter points showing relative DSB frequency versus distance to the nearest R-loop summit. DSBs were significantly enriched within 4 kb of R-loops in untreated cells, determined by a Wilcoxon test, adjusted p values: U2-OS = 7 × 10⁻⁷; HeLa = 1.6 × 10⁻⁵). This Enrichment persisted but was 2-fold weaker under aphidicolin (U2-OS = 7 × 10⁻⁷; HeLa = 5.5 × 10⁻⁸).

**Figure 3.**
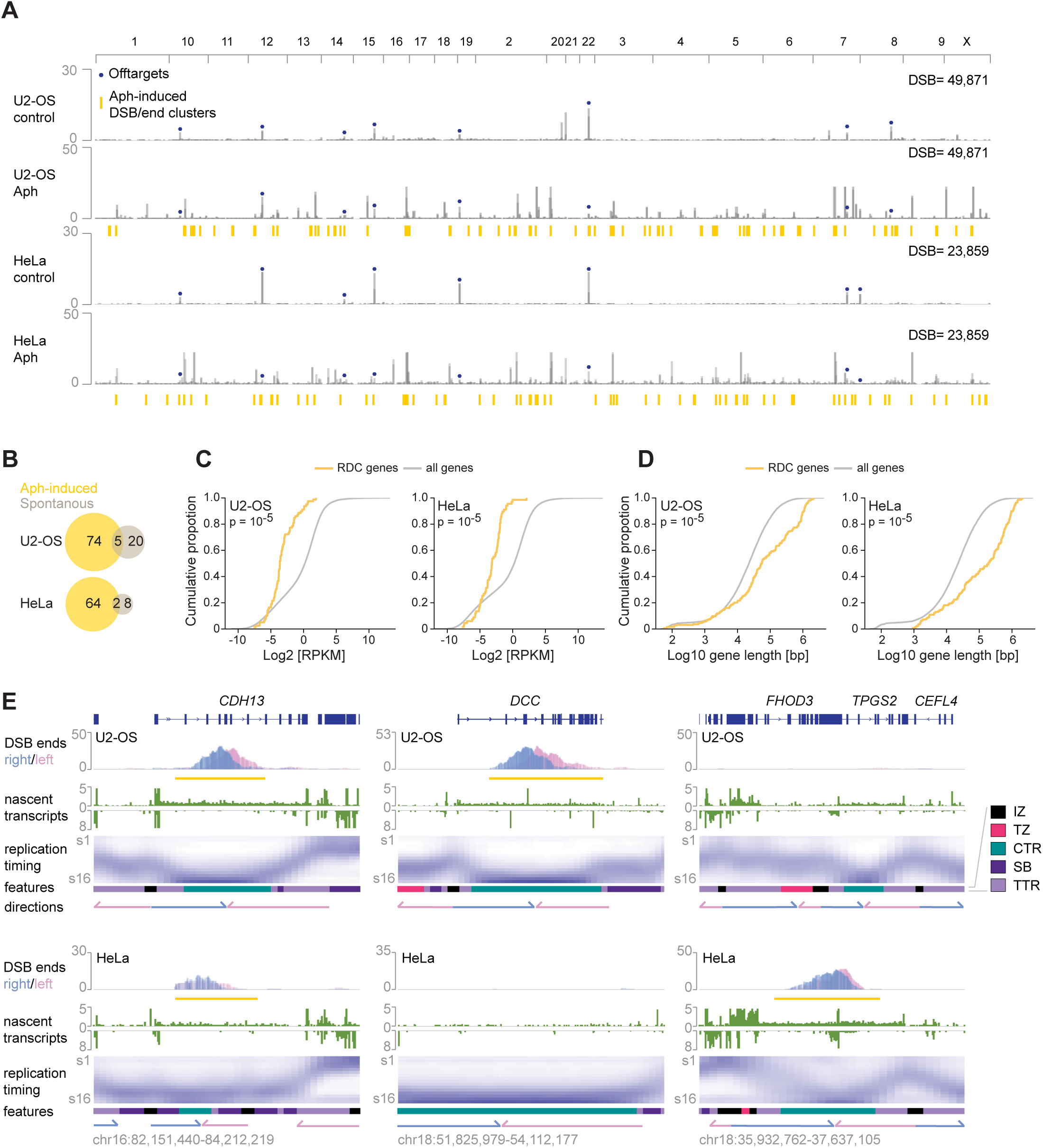
Mapping and genomic features of RDCs under replication stress. **(A)** Genome-wide mapping of RDC clusters in U2-OS and HeLa cells. Single-ended DSB densities were shown for DSBs with either a leftward end (pink) or a rightward end (blue). Significant Aph-inducible DSB clusters are annotated as orange bars below the Aph tracts. DSB clusters colocalized at Cas9 off-target sites are annotated with blue dots. **(B)** Venn diagram showing the intersection between RDC clusters detected under unperturbed conditions (intrinsic) and those induced by Aph treatment in U2 - OS and HeLa cells. **(C)** Empirical cumulative distribution functions (ECDFs) of gene expression for RDC genes (yellow) and all genes (grey) in U2-OS and HeLa cells. Gene sets matching the RDC set size were randomly sampled 100,000 times from all genes, and observed mean RDC expression was compared with this null distribution using a two-sided permutation test. Permutation p-values are shown. **(D)** Empirical cumulative distribution functions (ECDFs) of gene length for RDC genes (yellow) and all genes (grey) in U2-OS and HeLa cells. Gene sets matching the RDC set size were randomly sampled 100,000 times from all genes, and observed mean RDC gene length was compared with this null distribution using a two-sided permutation test. Permutation p-values are shown. **(E)** Multiomics figures provide information regarding genomic loci containing RDCs at the *CDH13*, *DCC*, and *FHOD* gene loci. The top panel shows the annotated gene structure. The second panel presents DNA break density mapped by LAM-HTGTS in U2-OS and HeLa cells, with pink and blue peaks representing leftward and rightward DSB ends, respectively. The third panel displays nascent transcript levels, reflecting transcriptional activity across the locus. The fourth panel depicts replication timing as a HiRepli-seq heatmap. The bottom panel provides domain annotations derived from replication timing, replication fork directions indicated by arrows, and additional genomic features.

**Figure 4.**
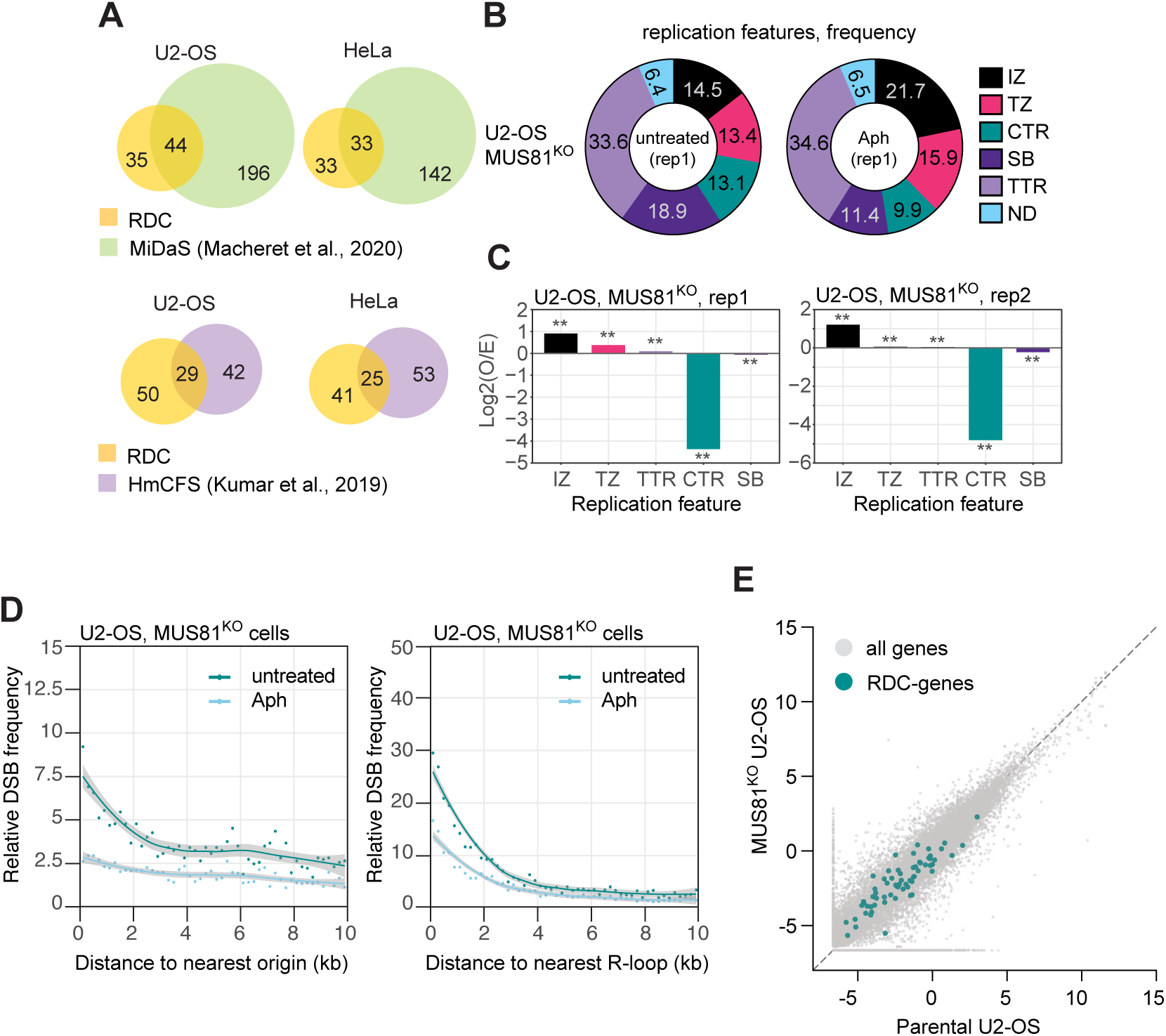
DSB, R-loop, and transcription landscapes in MUS81-deficient U2-OS and HeLa cells. **(A)** Venn diagrams showing overlap of RDCs with annotated RDC candidates, MiDAS sites and known human CFS (HmCFS) in U2-OS and HeLa cells. **(B)** Distribution of DSBs across replication features in untreated and aphidicolin (Aph)-treated MUS81^KO^ U2-OS cells (represented: replicate 1). Values indicate the percentage of DSBs falling within each feature: initiation zones (IZ), termination zones (TZ), constant timing regions (CTR), stochastic breaks (SB), timing transition regions (TTR), and regions with no assigned feature (ND). **(C)** Enrichment of R-loops at each replication feature relative to expectation for two independent replicates of MUS81^KO^ U2-OS cells. Bars represent log2 observed/expected score (Log2(O/E)); **p < 0.01 by permutation test. **(D)** Relative DSB frequency as a function of distance to the nearest replication origin (left) and nearest R-loop (right) in untreated and Aph-treated MUS81^KO^ U2-OS cells. Points represent binned observations; lines and shading indicate LOESS fit ± 95% confidence interval. **(E)** Gene expression in MUS81^KO^ versus parental U2-OS cells, measured by GRO-seq and expressed as log2 RPKM. Each point represents a gene; all genes are shown in grey and RDC-genes are highlighted in teal. The dashed line indicates equal expression between genotypes (linear regression, slope = 0.99).

**Figure 5.**
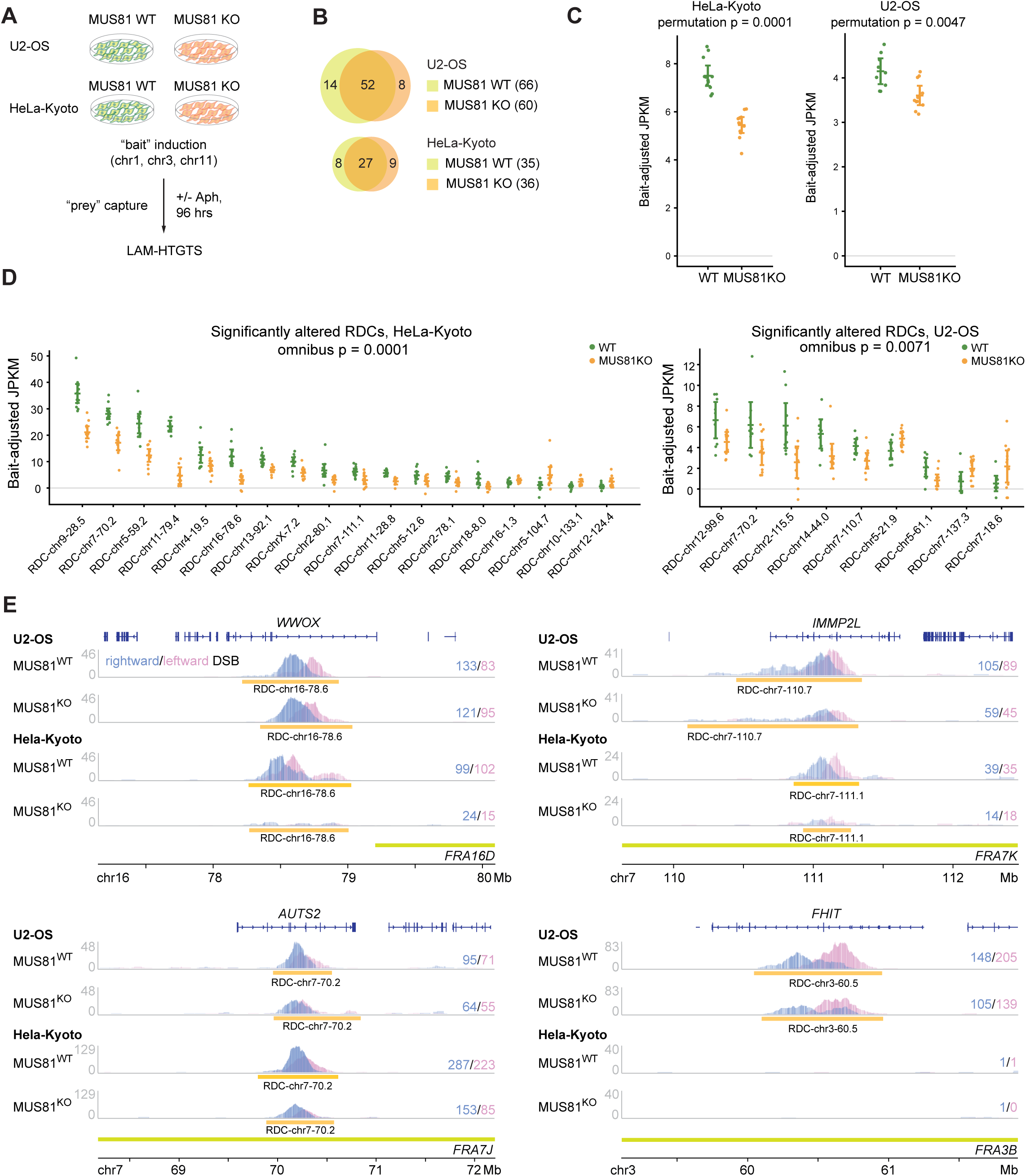
MUS81 dependency of RDC formation in U2-OS and HeLa-Kyoto cells. **(A)** Experimental design. MUS81^WT^ and MUS81^KO^ cells derived from U2-OS and HeLa-Kyoto parental lines were subjected to LAM-HTGTS with bait DSBs on chromosomes (chr) 1, 3, or 11, with or without aphidicolin (Aph) treatment for 96 hr. **(B)** Venn diagrams showing overlap of RDCs identified in MUS81^WT^ and MUS81^KO^ cells. **(C)** Mean junction density across all RDCs was compared between WT and MUS81^KO^ cells in U2-OS and HeLa-Kyoto. RDC junction counts were normalized to total junctions per library and RDC length and adjusted for chromosomal bait. Each point represents an individual capture library; horizontal bars and error bars indicate the mean and 95% confidence interval. Significance was assessed using a two-sided Welch statistic with permutation of genotype labels within bait groups. **(D)** Quantification of nominally significant (p<0.05, uncorrected) altered RDCs in U2-OS and HeLa-Kyoto. Each dot represents a biological replicate; error bars denote the mean with 95% confidence interval. Statistical significance was determined using a two-sided Welch statistic with permutation of genotype labels within bait groups. The significance level listed here comes from an omnibus permutation test of overall significance which provides weak control over the family-wise error rate (FWER). **(E)** Genome browser views of representative loci. Figure shows DSB density for MUS81^WT^ and MUS81^KO^ cells at the *WWOX*, *IMMP2L*, *AUTS2*, and *FHIT* loci. The corresponding fragile sites associated with genes are highlighted in lime green. Numbers to the right indicate single ended DSB counts per genotype.

### Assessing Changes in Replication Timing

To quantify aphidicolin-associated changes in replication-feature annotation while accounting for spatial dependence between neighboring Repli-seq bins, we performed a block-level paired transition analysis. Repli-seq feature calls were first represented on the 50-kb bin framework used for feature annotation, with each genomic bin assigned a feature label in untreated and aphidicolin-treated conditions. For each genomic block, we counted feature gains and losses for each annotation class, for example non-IZ → IZ versus IZ → non-IZ and calculated a net transition score as gain minus loss, normalized by the number of callable bins in the block.

Statistical significance was assessed using a paired chromosome-level sign-flip test. Briefly, for each feature and block size, aphidicolin-minus-untreated net transition scores were calculated independently for each chromosome. Under the null hypothesis of no consistent aphidicolin-associated gain or loss of a given feature, the sign of each chromosome-level difference is exchangeable. We therefore generated an empirical null distribution by randomly flipping the signs of chromosome-level transition scores and comparing the observed genome-wide mean transition score with this distribution. For experiments with two replicates, transition scores were calculated separately for each matched replicate pair and then combined at the chromosome level before significance testing.

### Comparing JPKM Values

RDC junction counts were normalized to total library junctions and RDC length to obtain JPKM. Group differences were assessed using Welch’s *t*-statistic with permutation-based significance testing. Where multiple chromosomal baits were used, bait effects were regressed out, labels were permuted only within bait groups, bait-chromosome RDCs were excluded, and baits lacking both conditions were removed. The same permutation was applied across RDCs to preserve inter-locus correlations. For individual RDCs, an uncorrected, two-sided permutation *p*-value was calculated, with RDCs exhibiting an effect at the *p* < 0.05 uncorrected level shown in figures. This is a non-parametric test that does not require us to make any assumptions about the underlying data distribution. Family-level evidence was assessed using an omnibus permutation test based on the sum of squared RDC-specific Welch statistics, allowing heterogeneous effects in either direction. This kind of test provides weak control over the family-wise error rate. We do not make strong inferences about individual RDCs.

A complementary global test applied the same Welch permutation procedure to each library’s mean JPKM across the relevant RDCs. Thus, the global test detects a consistent overall shift, whereas the omnibus test detects heterogeneous locus-specific effects. Where RDCs were divided into predefined genomic classes, both tests were repeated within each class.

As a quality control step, when comparing JPKM values, we only included samples with at least 750 total breaks across the whole genome.

### Quantifying enrichment of DSBs in Different Replication Timing Regions

LAM-HTGTS junction intervals were represented by their midpoint and assigned to 50-kb genomic bins, matching the resolution of the Repli-seq annotations. Leftward and rightward timing-transition regions were combined into a single TTR category. DSB enrichment in CTR, IZ, TTR, TZ and SB regions was assessed separately for each cell line, treatment and Repli-seq replicate. Null distributions were generated by independently circularly shifting the complete binned DSB-count track along each chromosome, thereby preserving chromosome-specific DSB totals and the spatial clustering and autocorrelation of DSBs. Enrichment was expressed as the observed DSB count divided by the mean count from 10,000 permutations. A one-sided upper-tail p-value was calculated as (1+*number of permutations with a DSB count greater than or equal to the observed count* ) / ( *n_permutations*+1). Therefore, only positive enrichment was tested; depletion and other two-sided departures from the null were not tested. P-values were adjusted across all reported one-sided enrichment tests using the Benjamini–Hochberg procedure.

### R-loop enrichment analyses

To address whether R-loops are enriched at RDCs, CFSs, or across replication features, we tested the overlap between R-loop peaks and each region-of-interest set against a permutation null. R-loop peaks were called with MACS2 and each peak was reduced to its summit to give an unambiguous point for overlap assignment. We then counted how many R-loop summits fell within the RDC set and within the CFS set. To define the expectation under no association, we randomly repositioned the summits 1000 times within the mappable, non-blacklist genome, that is the assembly minus the ENCODE blacklist, and recounted the overlap at each permutation. Randomization was performed on the same background from which the observed peaks were drawn, and observed counts were conditioned on peaks lying inside that background, so observed and expected values are directly comparable. We restricted the null to this background rather than the whole genome because R-loops preferentially form in mappable, actively transcribed regions, and an unrestricted null would inflate apparent enrichment. Enrichment was expressed as observed divided by the mean of the null (reported as log2 fold change), and significance as an empirical two-sided p-value from the permutation distribution, adjusted across the two region sets by the Benjamini-Hochberg test.

## Code availability

LAM-HTGTS, GRO-seq, Repli-seq pipelines are available at GitHub (https://github.com/brainbreaks/). Customized R scripts processing LAM-HTGTS output data are deposited also at the Github (https://github.com/brainbreaks/HTGTS-TLX-processing).

## Data availability

Raw and processed sequencing files were deposited at the European Nucleotide Archive (ERP181270). Processed data were aligned to hg38 genome build.

## Results

### Study Design

To directly compare how replication stress shaped genome fragility in cancer cells, we mapped replication dynamics, transcription, and DNA double-strand breaks in two widely used cell lines – U2-OS (osteosarcoma) and HeLa (cervical carcinoma) (13, 20–24) (**Figure 1A**). We generated multiple high-resolution cell-line-matched sequencing datasets, including DNA replication by high-resolution Repli-seq (HiRepli-Seq); transcriptional activity and orientation by global run-on sequencing (GRO-seq); and DNA double-strand breaks with end orientation by linear-amplification–mediated high-throughput translocation sequencing (LAM-HTGTS). DNA:RNA hybrid location was defined by DRIP-seq datasets. We generated DRIP-seq libraries for U2-OS cells, and we borrowed the HeLa results from prior studies (25). In addition, we chose aphidicolin to induce replication stress, as it is the canonical agent used to induce CFSs, allowing a direct comparison to our findings. Lastly, the TP53 pathway is either mutated or inactivated in HeLa and U2-OS cells (26). As loss of TP53 promotes tolerance to replication stress, and may facilitate the accumulation of recurrent DSBs, this shared deficiency provides a relevant context for cancer cells deficient in TP53.

### Cell-type-dependent replication feature shifts induced by long-term exposure to mild replication stress

The first question we asked was whether replication stress differentially altered DNA replication features. DNA replication features – initiation zones (IZs), timing transition regions (TTRs), termination zones (TZs), constant-timing regions (CTRs), steady-breakage regions (SBs), and biphasic regions – are closely linked to genome fragility (**Figure S1A**). These features capture distinct aspects of replication architecture. IZs are regions of preferential replication initiation, whereas TTRs undergo a gradual change in replication timing and are typically replicated predominantly by a long-traveling unidirectional fork, progressing with limited local origin firing. TZs mark regions where converging forks terminate, while CTRs exhibit relatively uniform replication timing across extended genomic intervals and can occur at different positions in S phase. Thus, “late” and “constant” describe distinct properties: a late CTR is both late-replicating and relatively uniform in replication timing. In neural progenitor cells, long TTRs are prone to transcription–replication conflicts, whereas late CTRs and biphasic regions are strongly associated with common fragile sites (CFSs) (6). Finally, SBs exhibit a nearly flat replication-timing profile spanning hundreds of kilobases, consistent with dispersed origin firing during mid-S phase (27), but their response to replication stress remains poorly understood.

To investigate how replication stress influenced replication architecture and associated DNA break formation, we performed high-resolution Repli-Seq (HiRepli-Seq) to define the above-mentioned replication features (**Materials and Methods, Figures. 1B,C; Table S2**). This analysis revealed feature-specific remodeling upon aphidicolin treatment, with some replication features displaying greater plasticity than others.

In order to test whether aphidicolin-associated changes in replication-feature annotation were significant, we performed a block-level paired transition analysis (see Material and Methods). In U2-OS, aphidicolin caused a significant net gain of TTR annotations (+2.51 percentage points; 95% block-bootstrap CI: +1.76 to +3.27; FDR = 0.002), together with significant losses of CTRs (−1.76 percentage points; 95% CI: −2.74 to −0.78; FDR = 0.013) and SBs (−2.28 percentage points; 95% CI: −3.02 to −1.60; FDR = 0.002). IZs and TZs showed small net gains in U2-OS, but these were not significant in the two-replicate combined analysis. In HeLa, aphidicolin instead produced a strong net gain of IZs (+5.23 percentage points; 95% CI: +4.10 to +6.35; FDR = 0.002) and a smaller but significant gain of SBs (+0.98 percentage points; 95% CI: +0.35 to +1.55; FDR = 0.019), accompanied by significant losses of TTRs (−3.06 percentage points; 95% CI: −4.00 to −2.27; FDR = 0.002) and CTRs (−2.44 percentage points; 95% CI: −2.92 to −1.94; FDR = 0.002). TZs showed a modest trend toward loss in HeLa, but this did not reach significance. Together, these results support cell-line-specific remodeling of replication-feature architecture after aphidicolin treatment, with U2-OS showing a shift toward TTR gain and SB/CTR loss, whereas HeLa shows prominent IZ gain with concomitant TTR/CTR loss.

The reduction of CTRs was somewhat counter intuitive as CTRs are linked to CFSs. Comparing locus-specific changes for CTR status revealed that mild replication stress selectively spared CFS-associated CTRs (28). CFS-associated loci including *WWOX*, *AUTS2* and *IMMP2L* genes, CTRs remained stable (**Figures 1D, E**). In summary, mild replication stress remodelled replication features in a cell-type-dependent manner. Steady breakages, whose stress response had not previously been characterized, changed significantly in both cell lines, but in opposite directions. Furthermore, CTR loss was not enriched at common fragile sites.

### Initiation Zones and R-loop-prone Genomic Regions Are Endogenous DSB Hotspots

To examine how DNA replication features related to DSB formation, we quantified DSBs captured by LAM-HTGTS (17) (**Figures 2A**). This assay uses a CRISPR/Cas9-induced “bait” DNA break to enrich for endogenous “prey” breaks and resolves the orientation of captured ends at the nucleotide resolution, providing insights into fork remodeling upon stalling. We first examined whether DSBs scaled with the genomic proportion of replication features. In untreated cells, more than a quarter of all DSBs localized to initiation zones (IZs), far exceeding their genomic coverage and indicating significant enrichment (p < 0.00005, permutation test, **Figure 2B**). IZs were also enriched for conserved origins (**Figure S1B,C**). Other replication features did not show significant enrichment for DSBs. On the contrary, in aphidicolin-treated cells, breaks accumulated predominantly in late-replicating CTRs (p < 0.00005, permutation test, **Figure 2B**), indicating that long term exposure to mild DNA polymerase inhibition selectively sensitizes origin-poor domains to breakage.

Because IZs were expected to undergo origin-firing conflicts, we next measured the distance from each DSB to the nearest human conserved origin (29) (**Figure 2C**). In untreated cells, DSBs peaked within ∼2 kb of origin, but this enrichment dropped after aphidicolin treatment, consistent with a shift toward breaks in origin-sparse CTRs. We then asked whether R-loop–prone regions contribute to the observed enrichment. We found that R loops were overrepresented in IZs and depleted in CTRs in both U2-OS and HeLa cells (**Figure S1D**). Consistent to this finding, DSBs were concentrated within four kilobases of R-loop summits in untreated cells (**Figure 2D**). This enrichment persisted but was weaker under aphidicolin, again consistent with increased breakage in R-loop-poor regions. In summary, DSBs were enriched at IZs, and chronic, mild replication stress created significantly more DSBs at the late CTRs.

Together, these analyses represented two distinct modes of genome instability. In the absence of exogenous stress, endogenous DSBs accumulated at origin-rich IZs that also harbored R-loop– prone regions. By contrast, inhibition of DNA polymerase activity shifted DSB formation to late-replicating, origin-poor regions that were not susceptible to R-loops.

### Aphidicolin Induces Recurrent DSB Clusters In Late Constant Timing Regions

These observations raised the question of whether the dispersed breaks observed under stress converged into large, recurrent clusters resembling CFSs or mitotic DNA synthesis (MiDAS) regions—both of which require DSB intermediates (13, 20, 30–32). In order to characterize the genomic location of DSBs enrichment, we mapped and classified recurrent DNA break hotspots genome-wide in U2-OS and HeLa cells exposed to long-term, low-dose aphidicolin.

We applied an established DSB clustering algorithm (6) to identify recurrent DNA break cluster (RDC) hotspots with orientation of captured DSB ends (**Figure S2A**). We applied this method to U2-OS and HeLa cells treated with low-dose aphidicolin for a long period, 96 hours (**Figure 3A** and **Table S4**). In untreated cells, only a few hotspots were detected, as most DSBs were widely dispersed across the genome. Some were attributable to off-target activity of the CRISPR/Cas9 used to generate bait, while others, such as those near *PVT1*, *BRF1*, and *HSP90AA1*, overlapped R-loop-enriched regions (**Figure S2B**). In contrast, low-dose aphidicolin induced 79 and 66 RDC hotspots in U2-OS and HeLa cells, respectively, with little overlap with untreated cells (**Figure 3B**). Most RDC hotspots were mapped to late-replicating constant timing regions, only a few occurred in timing transition regions (**Table S4**). RDC hotspot overlapped with genes or actively transcribed, unannotated genomes. These regions were typically weakly transcribed (**Figure 3C**) and substantially longer than average active genes (**Figure 3D**), resembling CFS-like RDCs initially defined in neural progenitors (6). We further validated our findings using an independent DNA break mapping assay, sBLISS in U2-OS cells. Consistent with the LAM-HTGTS results, sBLISS revealed that DNA breaks were enriched within RDC-containing genomic regions (**Figure S2C-E**).

In summary, we demonstrated that long term exposure to mild replication stress in cancer cells generated discrete RDCs concentrated in long, late-replicating genes rather than a diffuse increase in breakage across the genome.

LAM-HTGTS also revealed the orientation of captured DSB ends, reflecting replication-fork direction at the timing of stalling. At genomic loci where two opposite replication directions meet, DSB clusters display a bidirectional signature. This signature, which has been linked to error-prone repair and copy-number loss at RDCs in neural progenitors (33), was likewise evident in cancer cells. For example, at the *CDH13* gene locus, two opposing DSB peaks indicated forks arriving from a nearby IZ and from upstream steady-firing origins outside canonical IZs (**Figure 3E**). A similar configuration at the *SOX5* locus (**Figure S2F**) showed partially overlapping peaks, resembling those found in CTR-like RDCs in neural progenitors.

A substantial subset of RDCs displayed cell-line specificity (**Figure S2F**). U2-OS-specific hotspots occurred at *DCC* (**Figure 3E**), *SOX5*, and *MGAT4C* (**Figure S3F**), while HeLa-specific hotspots included *FHOD3* and *INPP4B* (**Figures 3E, Figure S2G**). The specificity could be all explained by the replication feature shift, as most RDCs overlapped with large, late-replicating CTRs in both cell types. Instead, differences in transcription correlated with hotspot presence. At *DCC*, replication was delayed in both cell types, but the hotspot was absent in HeLa, which lacked transcription. Conversely, at *FHOD3*, the absence of a hotspot in U2-OS was explained by early replication timing and transcriptional inactivity. These data indicate that late replication creates a permissive environment for RDC formation, while transcription modulates their cell-type specificity.

### MUS81-deficient cells significantly altered replication features in U2-OS cells

Although RDCs and M-phase abnormalities both landed on transcribed genes in late replicating domains, we found that only a subset of DSB hotspots overlapped with regions prone to MiDAS or CFSs (**Figure 4A**). This prompted us to ask whether RDCs arose through the same mechanisms that underlie MiDAS and CFS formation (22, 34). To this end, we investigated the role of MUS81 and GEN1, structure-specific endonucleases preventing M-phase genome aberrant (13, 35, 36), in RDC formation. While both MUS81 and GEN1 recognize branched DNA structures and have established roles in resolving replication-associated intermediates (30, 32, 37, 38), their involvement in RDC formation has not been investigated.

First, we tested whether MUS81 loss was associated with net changes in replication-feature annotation. We performed a paired transition analysis using whole chromosomes as permutation and bootstrap blocks for U2-OS cells expressing MUS81 (**Figure 1C**) and deficient in MUS81 (MUS81^KO^, **Figure 4B**) (also see Material and Methods). In untreated U2-OS cells, MUS81^KO^ produced a strong net gain of SB annotations (+4.55 percentage points; 95% chromosome-block bootstrap CI: +3.85 to +5.32; FDR = 0.002), together with significant losses of IZs (−3.69 percentage points; 95% CI: −4.65 to −2.87; FDR = 0.002) and CTRs (−1.24 percentage points; 95% CI: −1.99 to −0.54; FDR = 0.002). The small gains in TTRs and TZs were not significant in the two-replicate combined analysis. In aphidicolin-treated U2-OS cells, MUS81^KO^ instead produced a significant net gain of IZs (+1.43 percentage points; 95% CI: +0.73 to +2.18; FDR = 0.009) and a significant loss of CTRs (−2.11 percentage points; 95% CI: −2.57 to −1.70; FDR = 0.006).

We followed-up on this analysis by mapping DSBs **(Figure 4C)**, transcription profiles, and R-loops in MUS81^KO^ U2-OS cells. We found that MUS81 deficiency did not affect R-loop distribution in relation to replication features (**Figure 4C**). Similarly, MUS81 deficiency did not alter DSB enrichment relative to replication origins or R-loops (**Figure 4D**), nor did it affect transcriptional activity at genes containing RDCs (**Figure 4E**).

Together, these comparisons reveal distinct MUS81^KO^-associated patterns in untreated and aphidicolin-treated cells: untreated cells showed prominent SB gain with IZ/CTR loss, whereas under aphidicolin, the significant changes were restricted to IZ gain and CTR loss.

### DSB density in RDC reduced in MUS81-deficient U2-OS and HeLa cells

Next, we asked whether MUS81 deficiency altered RDC formation (**Figure 5A**). Here, we included HeLa-Kyoto cells (14), an independent lab-derived subclone of HeLa, in our assays. To avoid confusion with cells used earlier in this article (**Figures 1-3**), subsequent analyses of RDC changes under MUS81-deficient conditions were restricted to matched parental (MUS81^WT^) – offspring (MUS81^KO^) pairs. To avoid confusion with cells used earlier in this article (**Figures 1-3**), subsequent analyses of RDC changes under MUS81-deficient conditions were restricted to matched parental (MUS81^WT^) and offspring (MUS81^KO^) pairs. We also generated R-loop maps (DRIP-seq), transcription profiles (GRO-seq), and high-resolution Repli-seq profiles from MUS81^KO^ U2-OS cells.

Under mild aphidicolin treatment, we identified around 60 and 36 RDCs in U2-OS or HeLa-Kyoto cells with MUS81KO, respectively (**Figure 5A,B**), many of these RDCs overlapped with those identified in other lab clones reported in earlier figures. In order to compare JPKM at RDCs between WT and KO, we aggregated RDCs found in MUS81KO, WT and GEN1KO (in U2-OS, see later in the text), removing duplicate (overlapping) RDCs. We then calculated whether MUS81 loss produced a broader change in RDC activity beyond effects at individual loci by comparing mean junction density across the complete RDC set. MUS81^KO^ cells showed a significant reduction in mean bait-adjusted RDC junction density in both U2-OS (*p* = 0.0047) and HeLa-Kyoto (*p* = 1 x 10⁻⁴) cells (**Figure 5C**). We next examined the effects of MUS81 loss at individual RDCs (**Figure 5D**). In U2-OS cells, nine RDCs showed nominally significant differences between WT and MUS81^KO^ cells, with the majority displaying reduced junction density following MUS81 loss, although increases were observed at a subset of loci. The effect was more widespread in HeLa-Kyoto cells, where 18 RDCs were significantly altered and most showed substantially lower junction density in MUS81^KO^ cells. Consistent with these locus-specific changes, omnibus permutation testing showed significant evidence for an effect across the RDC set in both U2-OS (*p* = 0.0071) and HeLa-Kyoto (*p* = 1 x 10⁻⁴) cells. Thus, the locus-specific reductions observed following MUS81 loss are accompanied by an overall decrease in RDC-associated junction formation.

Despite this general trend, U2-OS and HeLa-Kyoto cells showed locus-dependencies on MUS81. For instance, the RDC at *WWOX* was not affected in U2-OS cells, while it was significantly reduced in HeLa-Kyoto cells. In addition, RDCs were rarely abolished in U2-OS cells, whereas several loci lost nearly all DSBs in HeLa-Kyoto (**Figure 5E**).

To confirm that MUS81 deficiency did not bias the capture assay itself, we examined microhomology usage, bait resection length and translocation frequency at the CRISPR/Cas offtarget sites (**Figures S3A,B**). None of these parameters differed between parental and MUS81^KO^ lines, indicating that loss of MUS81 did not alter DSB capture by LAM-HTGTS.

In summary, mapping DSB distributions in lineage-matched U2-OS and HeLa-Kyoto lines revealed that MUS81 deficiency markedly reduced DSB density within RDCs. These findings indicate that MUS81 acts partially upstream of RDC formation, likely through promoting fork cleavage.

### MUS81 endonuclease activity suppresses RDC in MUS81^KO^ cells

To investigate whether RDC-associated DSB density depended on the endonuclease activity of MUS81, we reconstituted MUS81-knockout U2-OS cells with either wild-type human MUS81 or a catalytically inactive mutant (D338A/D339A), each fused to EGFP at the N terminus (**Figure 6A, S4A**). We confirmed the loss of catalytic activity of the mutant *in vitro* (**Figure S4B,C**). To minimize toxicity associated with MUS81 overexpression, expression of both constructs was driven by a low-expression promoter. We further sorted cells expressing high/medium EGFP from cells with no or low EGFP expression. In EGFP-expressing cells, both variants were expressed at the level of endogenous MUS81 protein in parental cells (**Figure 6B**). We observed EGFP-MUS81 expression in the EGFP-low cells, albeit at a level much lower than in the parietal cells. To ensure that subsequent analyses included cells expressing MUS81 at the endogenous level, we sorted EGFP-positive cells to analyze DNA break and cell cycle profile.

**Figure 6.**
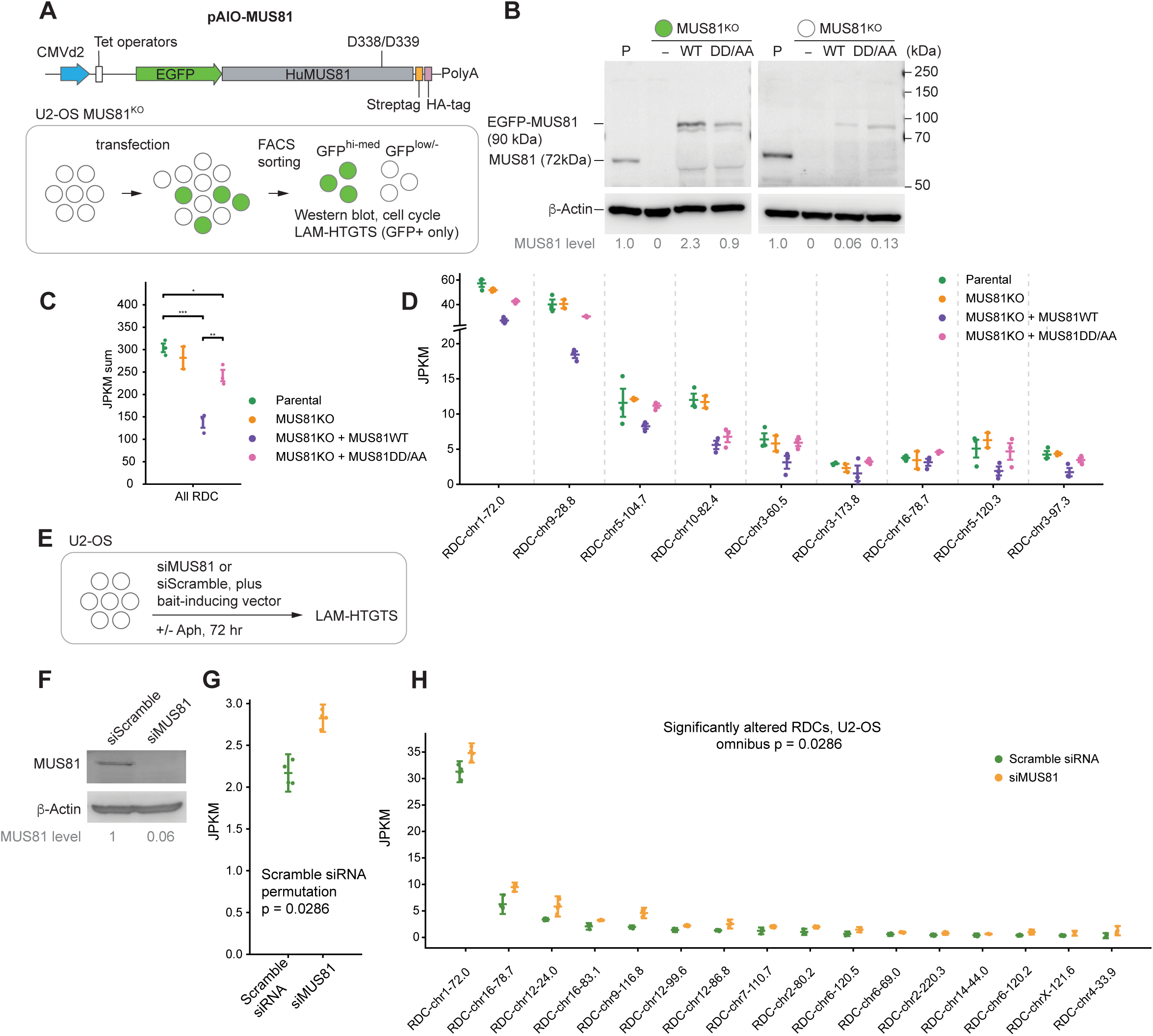
MUS81 catalytic activity was required to suppress RDC in MUS81^KO^ cells. **(A)** Top: the doxycycline-inducible pAIO-MUS81 construct. Bottom: U2-OS MUS81^KO^ cells were transfected, sorted by FACS into GFP+ and GFP-populations, and GFP+ cells used for western blot, cell-cycle analysis and LAM-HTGTS. **(B)** Western blot of parental (P), MUS81^KO^, and MUS81^KO^ cells reconstituted with MUS81^WT^ or MUS81^DD/AA^, comparing GFP+ (left) and GFP− (right) sorted fractions. The molecular weight of EGFP–MUS81 was approximately 28 kDa larger than endogenous MUS81 (61 kDa). β-actin, loading control. MUS81 levels normalised to β-actin and to parental cells are given below. **(C)** A comparison of the sum of JPKM values across all identified RDCs in parental U2-OS cells, MUS81-knockout cells (MUS81^KO^), and MUS81^KO^ cells reconstituted with wild-type MUS81 (MUS81^WT^) or the nuclease-dead mutant (MUS81^DD/AA^), determined by LAM-HTGTS from a chromosome 1 bait DSB. JPKM, junctions per kilobase per million junctions, normalized for RDC length and library size. **(D)** JPKM values for nominally significant (p<0.05 uncorrected) RDCs compared between different conditions using Welch’s two-sample t-test. Each symbol represents an independent library (n = 3 biological replicates per genotype); horizontal lines and error bars indicate mean ± 95 % confidence intervals. RDCs that show a significant (uncorrected) change between any of the conditions are displayed here RDCs. **(E)** Experimental scheme. U2-OS cells were transfected with siMUS81 or a non-targeting control siRNA (siScramble) together with the bait-inducing vector, treated ± aphidicolin (Aph) for 72 hr, and processed for LAM-HTGTS. **(F)** Immunoblot confirming MUS81 knockdown in siMUS81-relative to siScramble-transfected cells; β-actin, loading control. **(G)** Mean of JPKM values across all identified RDCs in siScramble- and siMUS81-treated cells. JPKM, junctions per kilobase per million junctions, normalized for RDC length and library size. **(H)** Quantification of nominally significant (p<0.05 uncorrected) RDCs between siScramble (green) and siMUS81 (orange). Each dot represents a biological replicate; error bars denote the mean with 95% confidence interval. Statistical significance was determined using a permutation test with a two-sided Welch test as the statistic of interest. The significance level listed here comes from an omnibus permutation test of overall significance which provides weak control over the family-wise error rate (FWER).

Reconstitution of MUS81 into MUS81-knockout U2-OS cells did not alter cell-cycle profile under the steady-state conditions (**Figure S4D,E**). Unexpectedly, it caused a pronounced S-phase arrest following prolonged treatment with long-term, low-dose aphidicolin, regardless of catalytic activity. We next performed LAM-HTGTS in the MUS81-reconstituted cells using a bait on chromosome 1 and compared the results with those from MUS81-knockout U2-OS cells processed in parallel. Reconstitution with wild-type MUS81 significantly reduced DSB density within RDCs to approximately 60% of that observed in parental cells (**Figures 6C,D**). In contrast, expression of the catalytically inactive mutant restored RDC-associated DSB density to the level close to that in MUS81-knockout cells.

These findings indicate that the catalytic activity of MUS81 is required to suppress RDC formation. If MUS81 were directly generating RDCs through cleavage of replication intermediates at the end of S phase, expression of the nuclease-dead mutant would be expected to reduce RDC-associated DSBs rather than restore them to the knockout level. Instead, our results support a model in which MUS81 protects genome integrity by catalytically resolving replication-associated DNA intermediates during early and mid S phase, thereby preventing their persistence into late-replicating regions where RDCs arise.

To further test this model, we transiently depleted MUS81 in MUS81-proficient U2-OS cells using siRNA (**Figures 6E,F**). As predicted, MUS81 knockdown increased DSB density within RDCs by approximately 25% compared with cells transfected with a scrambled siRNA control (**Figure 6G,H**). To our knowledge, this represents the first genome-wide segregation-of-function analysis demonstrating the contribution of MUS81 catalytic activity to DNA break formation. Collectively, these findings indicate that the endonuclease activity of MUS81 is required to resolve replication stress-associated DNA intermediates, thereby indirectly preventing RDC formation within late constant timing regions.

### GEN1 broadly inhibited while selectively promoted RDC formation in U2-OS cells

Next, we asked whether GEN1 contributed to RDC formation in U2-OS cells. We performed the same comparison in GEN1 knockout (GEN1^KO^) U2-OS cells, which shared the same parental origin as the MUS81^KO^ line (**Figure 7A**). Analysis of microhomology usage, bait length distribution, and off-target capture efficiency revealed that the loss of GEN1 did not impact DSB recovery by LAM-HTGTS (**Figure S5A-C**). By comparing mean junction density across the complete RDC set, we found that GEN1^KO^ cells showed a significant induction in mean bait-adjusted RDC junction density in U2-OS cells (permutation test, p = 0.0048) (**Figure 7B**). However, unlike in MUS81^KO^ cells, per-locus comparisons revealed that GEN1 played a dual role in RDC formation. A bait-adjusted omnibus test confirmed that 24 RDC loci showed a significant gain, whereas 4 (RDC-chr14-47.3, RDC-chr7-70.2, RDC-chr14-44.0, RDC-chr5-61.1) showed a significant loss, of DSBs in RDC (**Figure 7C**). For example, at the *DCDC1* and PARD3B loci,GEN1 loss promoted DSB accumulation (**Figure 7D**). Conversely, RDCs at *MDGA2* gene and an intergenic, actively transcribed genomic loci were significantly reduced in GEN1^KO^ cells (**Figure 7E**). As these loci all resided in late CTRs, differences were replication feature independent. In summary, unlike MUS81, which broadly promoted RDC formation, GEN1 modulated only a subset of RDCs.

**Figure 7.**
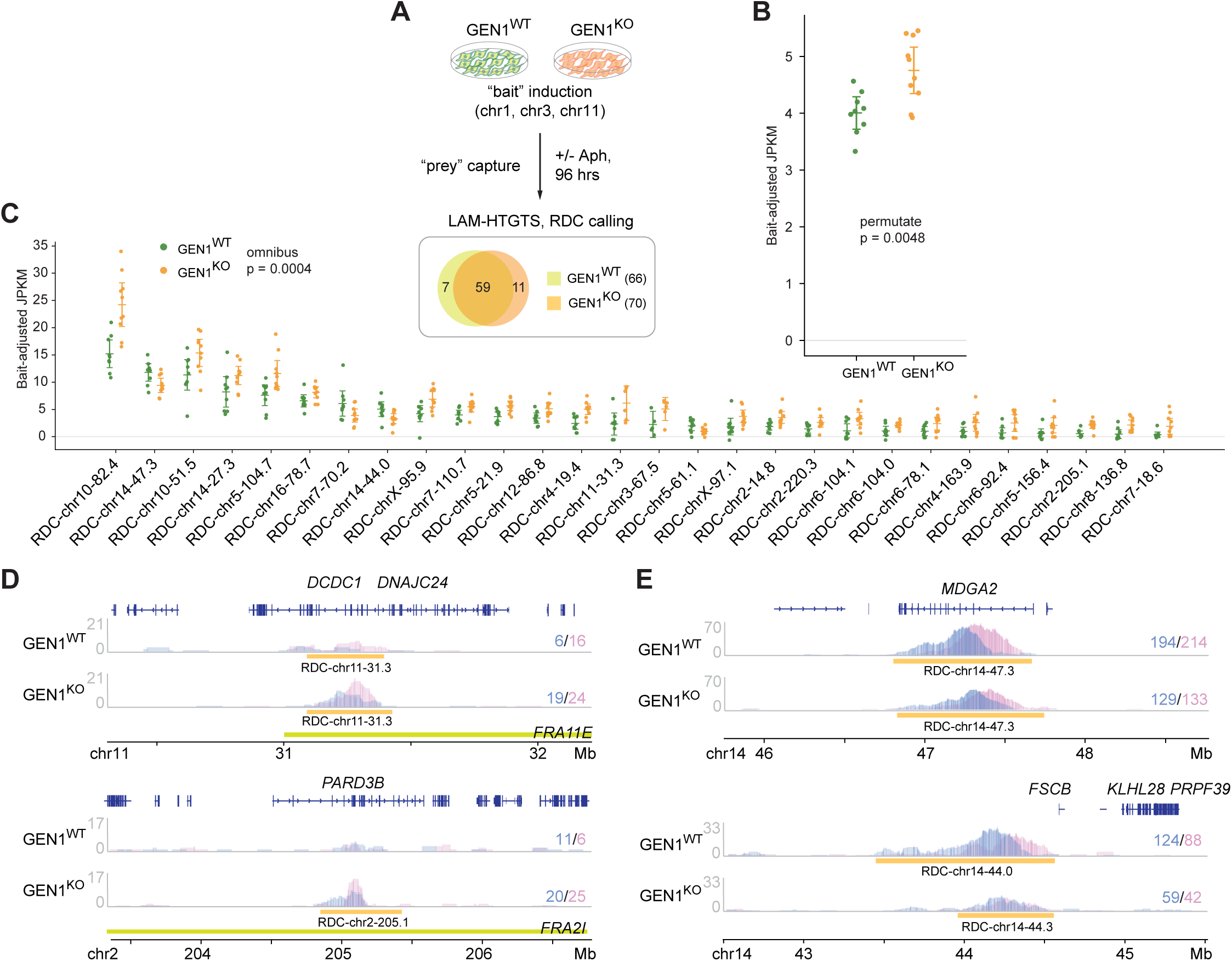
Locus-specific effects of GEN1 on RDC formation. **(A)** Schematic of experimental design. GEN1^WT^ and GEN1^KO^ U2-OS cells were subjected to LAM-HTGTS with bait DSBs on chromosome (chr) 1, 3, or 11, with or without aphidicolin (Aph) treatment for 96 hr. Venn diagram shows the overlap of RDCs identified in GEN1^WT^ and GEN1^KO^ cells. **(B)** Mean junction density across all RDCs was compared between WT and MUS81^KO^ cells in U2-OS. Figure is presented as described in Figure 5C. Significance was assessed using a two-sided Welch statistic with permutation of genotype labels within bait groups. (C) Quantification of nominally significant (p<0.05 uncorrected) RDCs in GEN1^KO^ U2-OS. Figure is presented as described in Figure 5D. The significance level listed here comes from an omnibus permutation test of overall significance, which provides weak control over the family-wise error rate (FWER). **(D,E)** Genome browser views of representative loci where RDC density was increased **(D)** or reduced **(E)** in GEN1^KO^ cells. Rightward or leftward single-ended DSB densities are shown for GEN1^WT^ and GEN1^KO^ cells, with RDC coordinates indicated below. Overlapping human common fragile sites are annotated as an FRA annotation. Numbers to the right indicate single-ended DSB counts within the given RDC per condition.

### MUS81 and GEN1 Differentially Regulate RDC-Associated DSBs Beyond Common Fragile Sites

Having established that MUS81 broadly promoted, and GEN1 selectively modulated individual RDCs, we compared both knockouts side by side to determine whether their effects follow common or distinct patterns across genomic contexts. In particular, we asked whether the loss of either nuclease preferentially affected fragile-site RDCs or extended to non-CFS regions.

A global comparison of loss versus gain across all RDCs revealed that MUS81 deficiency in U2-OS cells predominantly resulted in DSB loss at RDCs overlapping with CFSs, whereas DSB loss was marginal at RDCs not overlapping with CFSs (**Figure 8A**). In contrast, MUS81 knockout in HeLa-Kyoto cells resulted in an overall DSB loss regardless of CFS overlap (**Figure 8B**). On the other hand, GEN1^KO^ cells expressed a modest increase in DSB density at CFS-overlapping RDCs, whereas the increase was more pronounced at RDCs not overlapping with CFSs. Together, these results indicate that the differential effects of MUS81 and GEN1 loss on RDC-associated DSBs are not restricted to CFS-overlapping RDCs, suggesting that CFS overlap alone does not fully account for the differential sensitivity of RDCs to MUS81 or GEN1 deficiency.

**Figure 8.**
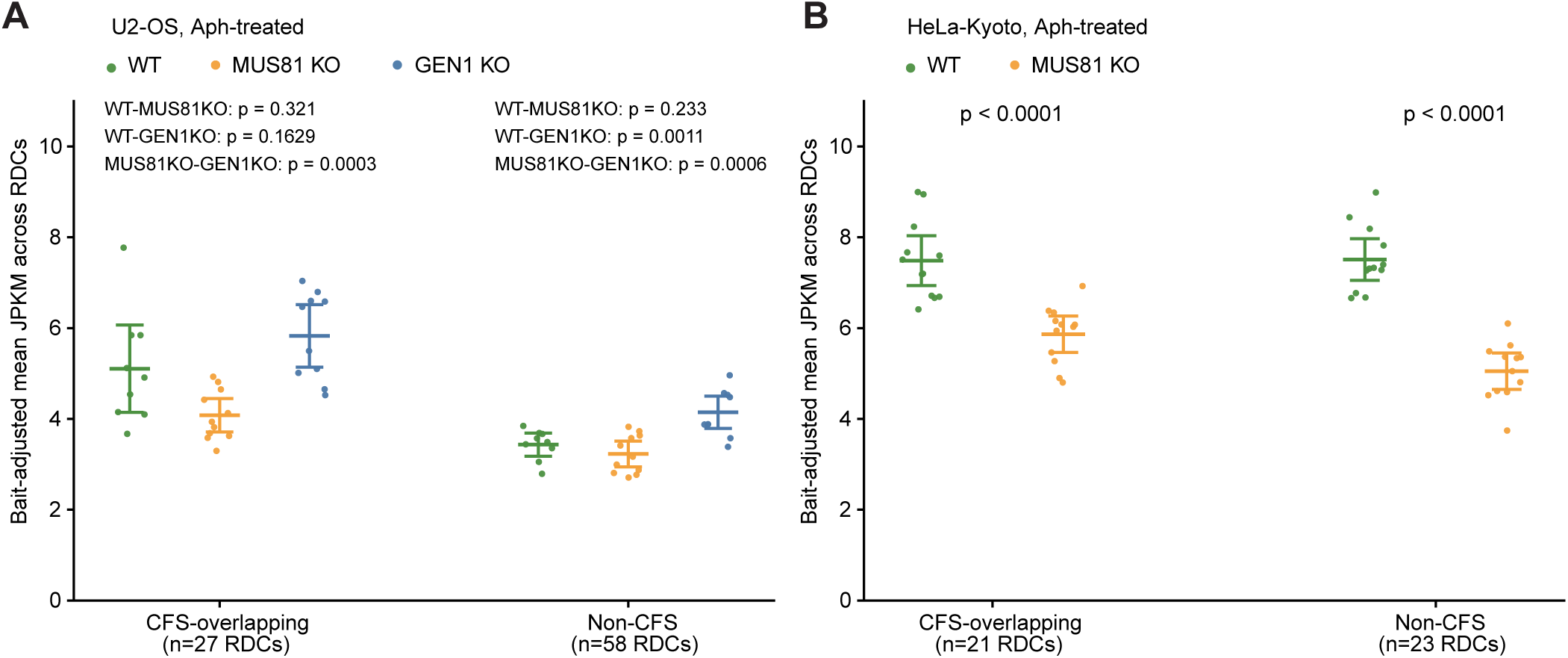
RDC dependency on MUS81 and GEN1 at CFS and non-CFS genomic regions. **(A)** Bait-adjusted mean JPKM across CFS-overlapping and non-CFS RDCs in aphidicolin-treated U2-OS WT, MUS81KO and GEN1KO cells. Each point represents the mean JPKM for one library across RDCs in the indicated class. Horizontal bars indicate group means and error bars show 95% confidence intervals. Bait-stratified permutation tests were performed between WT and MUS81KO, WT and GEN1KO, and MUS81KO and GEN1KO, separately within CFS-overlapping and non-CFS RDCs. **(B)** Bait-adjusted mean JPKM across RDCs overlapping common fragile sites (CFSs) and RDCs not overlapping CFSs in aphidicolin-treated HeLa-Kyoto WT and MUS81KO cells. Each point represents the mean JPKM for one library across RDCs in the indicated class; horizontal bars show group means and error bars indicate 95% confidence intervals. Genotype differences were assessed using a bait-stratified permutation test of the mean across RDCs. Genotype labels were permuted within bait groups, with the same permutation applied across RDCs.

## Discussion

### Novelty of the study

Our study provides a direct, genome-wide assessment of how replication stress reshapes the DSB landscape within the same cellular systems. By integrating LAM-HTGTS-based DSB mapping with replication timing, replication initiation features, transcriptional activity, and R-loop profiles, we distinguished endogenous and replication stress-induced DSB formation without relying on downstream measures of genome instability or comparisons across different biological contexts. While previous studies have identified individual genomic features associated with replication-associated instability (22, 25, 39–45), few have directly compared endogenous and stress-induced DSB landscapes together with their replication, transcriptional, and R-loop contexts in the same experimental system. Our analysis reveals that these DSB classes occupy distinct genomic environments: endogenous DSBs preferentially arise in origin-rich, R-loop-associated regions, whereas replication stress promotes break formation in late-replicating, origin-poor regions.

Our multi-omics analysis further establishes a specific relationship between R-loops and replication features: R-loops were enriched at initiation zones but depleted in late constant-timing regions, and their modest enrichment at common fragile sites was largely attributable to the subset of common fragile sites overlapping initiation zones. Consistent with this pattern, R-loops were also depleted within RDCs. Their distribution was largely unchanged following MUS81 loss, supporting the conclusion that MUS81 affects DSB formation without substantially altering the underlying R-loop or replication landscape.

We further demonstrated that cancer cells harbor RDCs, extending previous observations that had been limited to neural progenitor cells. Integrating RDCs with replication feature clustering showed that, in cancer cells treated with long-term, low-dose aphidicolin, more than half of RDCs in U2-OS and HeLa localized within late CTRs enriched for long, transcriptionally active genes. These features resemble CFS-like RDCs described in neural progenitor cells (6). In addition, we found that MUS81-deficient cells significantly altered replication features in U2-OS cells. Finally, segregation-of-function analyses revealed a universal contribution of MUS81 to RDC formation, with GEN1 contributing to a subset of RDC. Together, these findings establish an integrated framework linking replication architecture, transcription, R-loops, and nuclease activity to replication stress-induced genome fragility.

### Replication features describe fork architecture beyond replication timing

Replication timing and replication features are two complementary descriptors derived from HiRepli-Seq data. Replication timing collapses each genomic bin into a relative timezone (early or late) in S phase, whereas replication features are defined by the shape of the 16-fraction profile across neighboring bins, capturing where replication forks initiate, how they travel, and where they converge. Some features share the same replication timing but are formed by distinct biology. For example, steady-breakage regions (SBs) are origin-rich genomic regions, whereas timing transition regions (TTRs) are origin-poor domains; although these feature classes can share the same replication timing profile, they represent fundamentally different replication architectures. The ability to resolve initiation, elongation, and termination as separable replication events is a key advantage of high-resolution HiRepli-Seq (27), and our data distinguish these two descriptors in three ways.

First, aphidicolin remodeled replication features in opposite directions in identically treated cell lines: TTR annotation increased in U2-OS but decreased in HeLa, with reciprocal changes in IZs and SBs (**Figure 1B,C**). Second, loss of CTR annotation was not determined by late replication. CFS-associated CTRs at *WWOX*, *AUTS2*, and *IMMP2L* were retained, only 27% of CFS-overlapping CTR bins lost their annotation, and genome-wide CTR loss was not enriched at CFSs (**Figure 1D,E**). Third, replication features harboring breaks differed between systems occupying the same broad temporal window. RDCs occurred predominantly in TTRs in p53-deficient neural progenitors (6), but in late CTRs in U2-OS and HeLa. Although both feature classes replicate late in S phase, they represent fundamentally different replication architectures: TTRs reflect sparse initiation and long, largely unidirectional forks (46), whereas late CTRs are plateau-like domains that contain internal initiation events.

Late replication is therefore permissive rather than instructive for fragility. Only 10.7% of late CTRs in U2-OS and 9.7% in HeLa harbored an RDC, and ablating transcription in neural progenitors abolished DSB formation while further delaying local replication (6). Although aphidicolin-induced changes in replication timing have been characterized previously (41), our study addresses a different question. By pairing HiRepli-Seq feature annotation with orientation-resolved DSB mapping in the same cells, we identified the replication configuration in which breaks arise, showing that replication architecture, rather than replication timing itself, is the feature most closely associated with genomic fragility.

### Transcriptional activity, not R-loops, in the late-replication regions permits RDC formation in cancer cells

RDCs occurred predominantly within late CTRs, indicating that replication timing predisposes these domains to fragility. However, timing alone is insufficient: only a minority of late CTRs overlapped with RDCs. This late fragility has been previously suggested to result from under-replication of large genes or from transcription–replication conflicts in CFS (34, 44, 46). Although we detected under-replication signals within the *WWOX*, *IMMP2L*, *AUTS2*; *FHIT* RDCs (**Figure S6A**), not all RDC hotspots were underreplicated. This interpretation is consistent with our recent findings from a parallel study in mouse neural progenitor cells, in which abolishing transcription of RDC-containing genes eliminated DSB signals despite further delaying local replication timing (6). Together, these observations suggest that CTRs provide a permissive context for RDC formation, while transcription–replication conflicts contribute directly to DSB generation. On the other hand, although R-loops were slightly enriched within CFS regions, we found that they were markedly depleted across RDC-containing genomic regions (**Figure S6B-C**), consistent with previous findings in neural progenitor cells (6). These observations suggest that the contribution of transcriptional activity to RDC formation did not involve an R-loop-dependent DSB mechanism.

We speculate that elongating RNA polymerase II may be critical for recruiting fork remodeling complexes. For example, elongating RNA polymerase II interacts with RECQL5, a DNA/RNA helicase required for transcription (47–49). RECQL5’s helicase activity and recruitment to replication forks are also essential for MUS81-dependent fork cleavage (20). Whether the association of RECQL5 with RNA polymerase II is specifically required for MUS81-dependent RDC formation remains to be tested. In parallel, transcription-coupled nucleotide excision repair can generate single-stranded gaps (50), which, upon frequent transcription-replication conflict-mediated fork stalling, may be converted into single-ended DSBs (51, 52) independent of MUS81. Determining whether these pathways account for the transcriptional dependency of RDC formation will require further investigation.

### Transient and long-term loss of MUS81 have opposite effects on replication stress-mediated DNA breaks

We demonstrated that most RDCs were MUS81-dependent, irrespective of their association with CFSs (Figures 4, 5, 7). MUS81 has previously been shown to process stalled replication intermediates and promote the replication stress response (36, 53). Our results further revealed that functional MUS81 protected cells from RDC formation, whereas the nuclease-dead MUS81 mutant did not (**Figure 6**). Transient depletion of MUS81 increased RDC levels, indicating that its catalytic activity is primarily required to resolve replication-associated DNA intermediates before cells enter late S phase. Notably, expression of the nuclease-dead MUS81 mutant reduced RDC levels by approximately 20%, suggesting that MUS81-mediated fork cleavage directly contributes to a subset of RDC-associated DSBs.

The finding that transient and chronic loss of MUS81 has opposite effects on DNA break induction under replication stress is puzzling. Here, we propose a possible explanation for this discrepancy. We hypothesize that MUS81-deficient cells initially accumulated elevated levels of unresolved replication intermediates and subsequently underwent a replication-stress crisis. By the time of our analysis, the surviving cells may have adapted by reprogramming their replication profiles, consistent with the significantly elevated steady-state breakage observed in these cells.(**Figure 4B**). Effectively, MUS81-deficient cells reduced the genomic area left to be replicated at the end of S phase, thereby indirectly reducing the chance of DNA breaks. In addition, we hypothesize that these replication intermediates can also be rerouted to other endonucleases that recognize the three-way junction at a stalled fork. Restoring functional MUS81 protein in these cells allows MUS81 to take over its role again, presumably in early-mid S phase, effectively reducing the persistence of replication intermediates into late S or G2/M phase. Transient silencing of MUS81, on the contrary, exposed cells to unresolved replication intermediates that persisted into late S or G2/M phase, which we believe were then converted to DNA breaks by GEN1 or other endonucleases processing these intermediates. Together, our findings demonstrate an important distinction between chronic and transient loss of MUS81, most likely arising from adaptation of DNA replication architecture, which should be taken into account in future studies of DNA damage.

Although unlikely, we cannot rule out that a subset of RDC arises from replication fork reversal. Upon fork stalling, the nascent strands anneal and the parental strands re-pair, remodeling the three-way fork into a four-way “chicken-foot” junction whose regressed arm is structurally equivalent to a one-ended DSB (55, 56). This arm can be incised by the structure-specific endonuclease MUS81 and its EME1/2 partners, typically downstream of MRE11/EXO1 resection, converting the reversed fork into a frank one-ended DSB channeled into POLD3-dependent break-induced replication (57, 58). Consistent with this route being only one of several, MUS81 depletion reduced RDC-associated breaks by ∼20% in our system, indicating that the majority (∼80%) of RDC breakage is independent from the nuclease activity of MUS81. Residual breakage is likely contributed by other structure-specific endonucleases acting on branched fork intermediates (e.g., SLX1–SLX4, XPF–ERCC1), by replication run-off at pre-existing single-strand nicks or gaps which generates a one-ended DSB without nucleolytic cleavage (57, 59). Once formed, resected ends may be further processed by RAD52-dependent single-strand annealing between flanking repeats, a RAD51-independent route yielding deletion-type rearrangements (60). In summary, these results separate the biogenesis of RDC from that of mitotic synthesis and CFSs.

### Mechanisms beyond MUS81 and GEN1 in RDC formation

RDCs arise from transcription–replication conflicts; therefore, transcription-coupled mechanisms are likely to contribute to their formation (33, 44, 61). For example, transcription-coupled nucleotide excision repair intermediates transiently leave a single-strand discontinuity (62). When these gaps are encountered by an approaching replication fork or persist into the next round of DNA synthesis, they can be converted into double-strand breaks (51, 52, 63). This gap-to-break mechanism has been demonstrated in both mammalian cells and yeast, primarily within early-replicating genomic regions. We speculate that a subset of DNA breaks in late-replicating RDCs may also arise through a similar mechanism.

In addition to transcription-coupled DNA damage, chromatin organization and nuclear architecture may also influence RDC formation. Our recent study in neural progenitor cells showed that most late-replicating RDCs localized within lamina-associated domains (64). Because the nuclear periphery experiences greater mechanical constraints than the nuclear interior, these regions may be subjected to elevated physical stress, increasing their susceptibility to DNA breakage under replication stress (65, 66).

Finally, RDCs may also arise after mitosis from unresolved replication intermediates. Under replication stress, incompletely replicated fragile loci remain interlinked as ultrafine anaphase bridges (67), which are normally resolved without breakage by the BTR complex acting with the PICH translocase (68). When resolution fails, bridges break during cytokinesis, and the resulting damage persists into the next G1 phase (69). A related route operates on the thicker chromatin bridges that survive into abscission, where the midbody-tethered nuclease ANKLE1 cleaves bridge DNA (70). Because bridge breakage deposits DNA ends in both daughter cells, each end can be joined only to a partner present in that same daughter, offering a route to structural variation that differs between sister cells.

## Limitations of the study

While aphidicolin is a potent tool, we acknowledge that part of our findings are specific to stress induced by aphidicolin-mediated DNA polymerase inhibition. LAM-HTGTS selectively captures DSBs or DNA ends that are translocation-ready. Single-strand breaks, chemically “dirty” DSBs, or breaks covalently bound to proteins or trapped in complexes are not detected. Likewise, DSBs repaired by homologous recombination - which represent a substantial fraction of breaks in S phase - are not captured. As a result, our analyses reflect relative rather than absolute changes in DSB levels.

## Acknowledgement

We thank the FACS core facility, the NGS core facility, and the ODCF data management core facility at the DKFZ. We thank Stephen West and Pavel Janscak for their generous cell line gifts (MUS81^KO^ and GEN1^KO^, as well as parental cell lines), and Rami Aqeilan for his critical feedback. We thank Marco Giaisi for managing the Wei lab. We thank Giulia Di Muzio, Hsin-Ju Lu, and Mila Duerink for providing technical support and resources for the project. We extend our acknowledgements to the Wel lab members for fruitful discussion and suggestions.

## Author contribution

B.D. and P.-C. W. conceptualized and designed the project. B.D., L. C., M.G., D. S., V.M., and P.-C. W. performed the experiments: HiRepliSeq (B.D. and M. G.), GRO-seq (B.D. and L.C.,), DRIP-Seq (P.-C. W.), biochemical assay (V.M), sBLISS (D.S.), and LAM-HTGTS (B.D., L.C., M.G.). A. I., B.D. and P.-C. W. analyzed the data and interpreted the results. A. I., B. D. and P.-C. W. created figures. A. I. , B.D., and P.-C. W. wrote the manuscript. R. A., L. K., and P.-C. W. supervised the experiments. P.-C. W. secured research budget, supervised the team, and oversaw the research.

## Supplementary Figure Legends

**Figure S1. Replication origins, R-loops, and transcriptional activity at replication features, related to Figure 2. (A)** A diagram showing replication features defined in HiRepli-seq results. IZ: initiation zone, TZ: termination zone, TTR: timing transition region, CTR: constant timing region, and SB: steady breakage. **(B)** Bar plot showing counts of conserved replication origins intersecting each replication feature in U2-OS cells. The data represent two HiRepliSeq replicates, the untreated condition. **(C)** Genome browser view of the *KCNK9–PTK2* locus. Human core replication origins (orange) are shown above the replication timing heatmap derived from HiRepli-Seq with untreated U2-OS cells. Replication features (IZ, TZ, CTR, SB, TTR) are annotated below. **(D)** R-loop enrichment is shown as observed/expected ratio in a log2 scale, log2(O/E), across five replication features: IZ, TZ, TTR, CTR, and SB. Observed counts are the number of R-loop peak summits assigned to each feature by maximal overlap. Expected counts are the mean over 1000 permutations in which summits were randomly repositioned within the mappable, non-blacklist genome. Positive values indicate enrichment, negative values depletion. Panels show two biological replicates each for U2 -OS (top) and HeLa (bottom). In all four datasets R-loops are enriched at IZ and depleted at TZ, TTR, CTR, and SB, with the strongest depletion at CTR. Significance is from the empirical two-sided permutation p-value, Benjamini-Hochberg adjusted: ** p_adj < 0.01.

**Figure S2. Cell-type–specific RDCs and their transcriptional correlates, related to Figure 3. (A)** Schematic showing single-ended DSB captured by LAM-HTGTS. At a pair of stalled replication forks, fork remodeling created one single-ended DSB at each replication stalling site. The DSB at the rightward-moving fork is born with a rightward orientation (blue), and DSB at the opposite fork inherit a leftward orientation (pink), respectively. DSB density reflecting the fork stalling directions are shown in LAM-HTGTS DSB density plots in Figure 3F and subsequent figures. **(B)** Multiomics figures provide information regarding genomic loci containing spontaneous RDCs in U2-OS and HeLa cells. The top panel shows the annotated gene structure. The second panel presents DNA break density, with pink and blue peaks representing leftward and rightward DSB ends, respectively. The third panel displays nascent transcript levels, reflecting transcriptional activity across the locus. The fourth panel shows R-loop formation measured by DNA:RNA hybrids. The bottom panel depicts replication timing as a Repli-seq heatmap. (**C-E**) sBLISS mapping of aphidicolin-induced DSBs in G2/M U2-OS cells identifies breakage at RDCs. **(C)** Experimental scheme. U2-OS cells were seeded (day 1), synchronized by thymidine block (day 2) and released (day 3). Two hours after release, cells were treated with 0.3 µM aphidicolin (Aph); after a further 4 h, the CDK1 inhibitor (CDK1i) was added, and cells were collected 14–16 h later (day 4) in G2/M for sBLISS library preparation. **(D)** Venn diagrams showing the number of annotated fragile regions recovered by the 852 sBLISS tiles significantly enriched for breaks upon Aph treatment (0.5 Mb tiles, DESeq2, FDR < 0.1; black). Colored circles denote U2-OS recurrent DSB clusters (RDC, n = 79; orange), mitotic DNA synthesis sites (MiDaS, n = 269; blue) and common fragile sites (CFS, n = 86; purple). Numbers indicate regions of each class that overlap at least one significant tile, and those that do not. Circle areas are not to scale. **(E)** Heat map of the significant sBLISS tiles that overlap a U2-OS RDC. Each row is one tile, labelled by genomic coordinate (left) and by the RDC it intersects (right). Columns show two independent sBLISS replicates for untreated (Aph −) and aphidicolin-treated (Aph +) samples. Color indicates the row-wise Z score of normalized break density (scale at right); rows are ordered by chromosome. Break signal at these loci is consistently low in untreated cells and elevated after replication stress. Paired Wilcoxon test comparing the Z scores in tiles shown in C suggests DSB density was significantly enriched in cells treated with Aph (p = 6.9 x 10-10). **(F,G)** Multiomics figures provide information regarding genomic loci containing U2-OS-specific RDCs **(F)** and HeLa-specific RDCs **(G)**. The top panel shows the annotated gene structure. The second panel presents DNA break density mapped by HTGTS in U2-OS and HeLa cells, with pink and blue peaks representing leftward and rightward DSB ends, respectively. The third panel displays nascent transcript levels, reflecting transcriptional activity across the locus. The fourth panel depicts replication timing as a Repli-seq heatmap. The bottom panel provides domain annotations derived from replication timing, replication fork directions indicated by arrows, and additional genomic features.

**Figure S3. MUS81 deficiency did not affect microhomology usage and end resection in U2-OS cells, related to Figure 5. (A)** Microhomology (MH) usage at junctions captured with chromosome (chr) 1, 3, or 11 baits in U2-OS (top) and HeLa-Kyoto (bottom) MUS81^WT^ and MUS81^KO^ cells. **(B)** Bait length (Blen) distributions for junctions captured with chr 1, 3, or 11 baits in U2 -OS (top) and HeLa-Kyoto (bottom). Shaded areas indicate expected full-length Blen peaks.

Figure S4. MUS81^WT^ and MUS81^DD/AA^ protein function in *in vitro* 3’-flap DNA cleavage and cell cycle progression in U2-OS cells, related to Figure 6. (A) Sanger sequencing result showing the coding sequences of the pAIO expression construct at D338/D339 of MUS81^WT^ (upper, GATGAC) and MUS81^DDAA^ mutant (lower, GCTGCC). (B) Biochemical characterization of MUS81 nuclease activity. Coomassie-stained SDS-PAGE of recombinant truncated wild-type (WT) and nuclease-dead (DD/AA) His-ΔEME1–ΔMUS81 complexes co-purified from E. coli. L, molecular weight marker (kDa); positions of truncated hisΔEME1 and ΔMUS81 are marked. (C) Nuclease assay on a labelled 3’-flap DNA substrate (asterisk denotes the labelled end) incubated with increasing concentrations WT (0– 2.5 nM) or DD/AA (0–5 nM) His-ΔEME1–ΔMUS81. Reaction products were resolved by native PAGE (top); the migration of uncleaved substrate and cleaved product is indicated schematically at left. The bar graph (bottom) shows quantification of cleavage by the WT complex (mean ± SD, n = 3). WT ΔMUS81–EME1 cleaved the substrate in a dose-dependent manner, whereas the DD/AA variant was catalytically inactive across all concentrations. **(D)** Representative DNA content histograms (propidium iodide) of the four cell lines, untreated (top) or treated with 0.4 µM aphidicolin (Aph, bottom). **(E)** Quantification of G1, S and G2/M fractions from (D) for untreated (grey) and Aph-treated (teal) cells. Mean ± SD of 3 independent experiments. Aph induced G1 accumulation in parental and MUS81^KO^ cells, which was partially reduced in both reconstituted lines. two-way ANOVA with cell line and treatment as fixed factors and experiment as a blocking factor (repeat n = 3) was conducted. Both log-ratios were significantly shifted (P < 0.0006), while in untreated cells, only the G2/M fraction differed between lines (P = 0.014), and no individual line differed from MUS81 ^KO^ after Dunnett correction. Statistical testing results were provided as Supplementary Table S1.

**Figure S5. GEN1 deficiency did not affect microhomology usage and end resection in U2-OS cells, related to Figure 7. (A)** Microhomology (MH) usage at junctions detected with chromosome (chr) 1, 3, or 11 baits in GEN1KO U2-OS cells. Frequencies are shown as a function of MH length (bp). **(B)** Bait length (Blen) distribution for junctions captured with chr 1, 3, or 11 baits. Shaded areas indicate corresponding full Blen peaks. **(C)** Off-target junction density (JPKM) in GEN1^WT^ and GEN1^KO^ cells. Note that the same dataset for GEN1^WT^ was shown as MUS81^WT^ in Figure S4D, as GEN1^KO^ and MUS81^KO^ cells share the same parental U2-OS line. No significant difference (n.s.) between two genotypes was observed.

**Figure S6. Under-replication and R-loop depletion properties at RDC loci in U2-OS cells, related to Discussion. (A)** Heatmaps show the under-replication index (URI) calculated from whole-genome sequencing data comparing aphidicolin (Aph)-treated U2OS cells to untreated controls. Four representative common fragile site genes (*WWOX, IMMP2L, AUTS2,* and *FHIT*) are shown. Two independent biological replicates are presented (rep1 and rep2, respectively). Genomic coordinates (hg38) are indicated below each locus. These data illustrate that APH treatment induces under-replication at multiple RDC-containing loci in U2-OS cells. **(B)** Permutation test of R-loop overlaps with RDC or CFS region set in U2-OS and HeLa cells. R-loop peak summits were randomly repositioned 1000 times within the mappable, non-blacklist genome and the overlap count was recorded per permutation. Blue bars, null distribution of overlap counts; red line, observed count from the real peaks. **(C)** Enrichment is summarized as log2(observed/expected), where expected is the mean of the null in (B). Positive values indicate enrichment, negative values depletion. RDC (blue) shows strong depletion (log2FE = -1.87) and CFS (yellow) a modest enrichment (log2FE = 0.11). Significance is from the empirical two-sided permutation p-value, Benjamini-Hochberg adjusted across the two tests: ** p-adj < 0.01.

## Code for data analysis

https://github.com/brainbreaks/

## Data availability

European Nucleotide Archive (ERP181270)

