## Supplementary figures and images for "An integrated genome-wide resource reveals distinct replication environments of DNA breakage in human cancer cell lines"

### Supplementary Figures 1-6

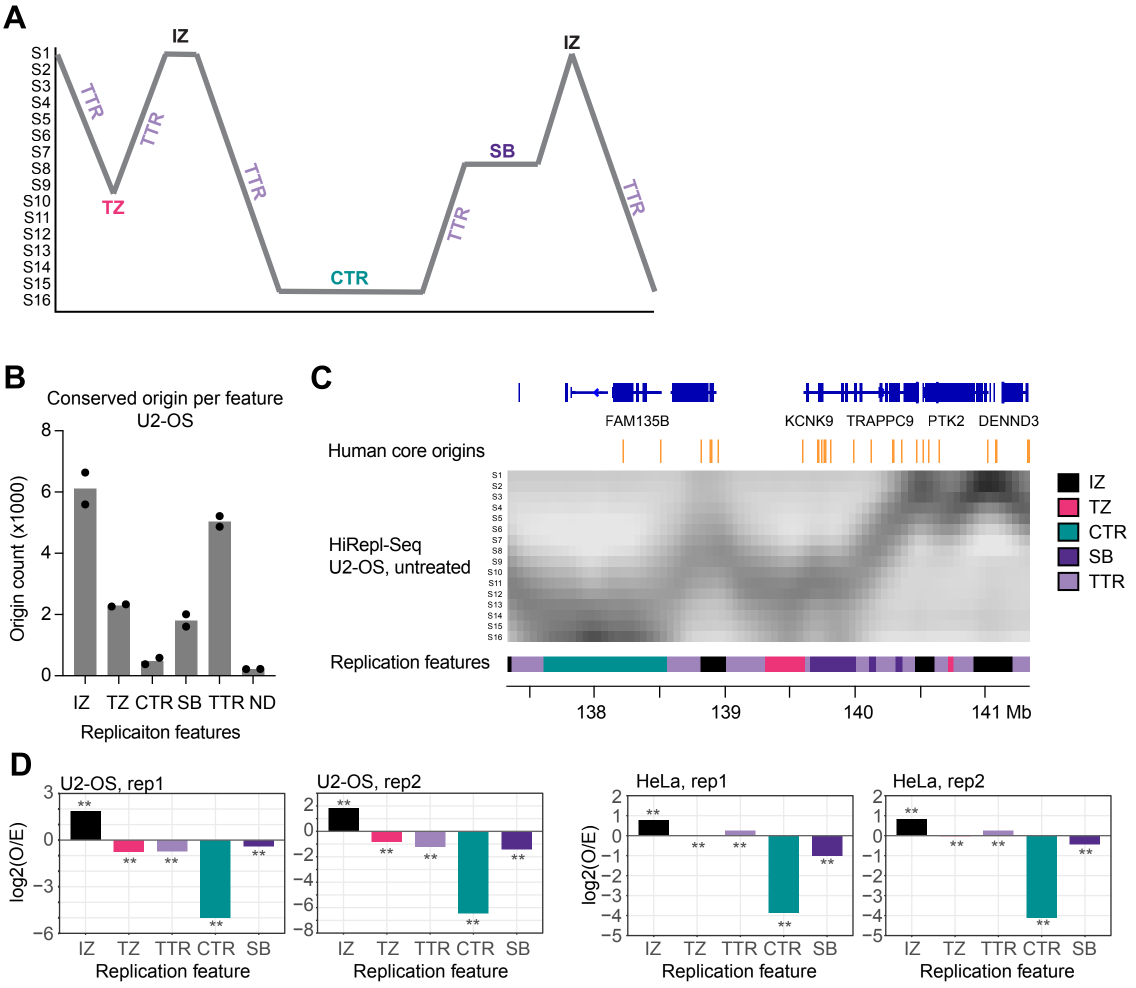

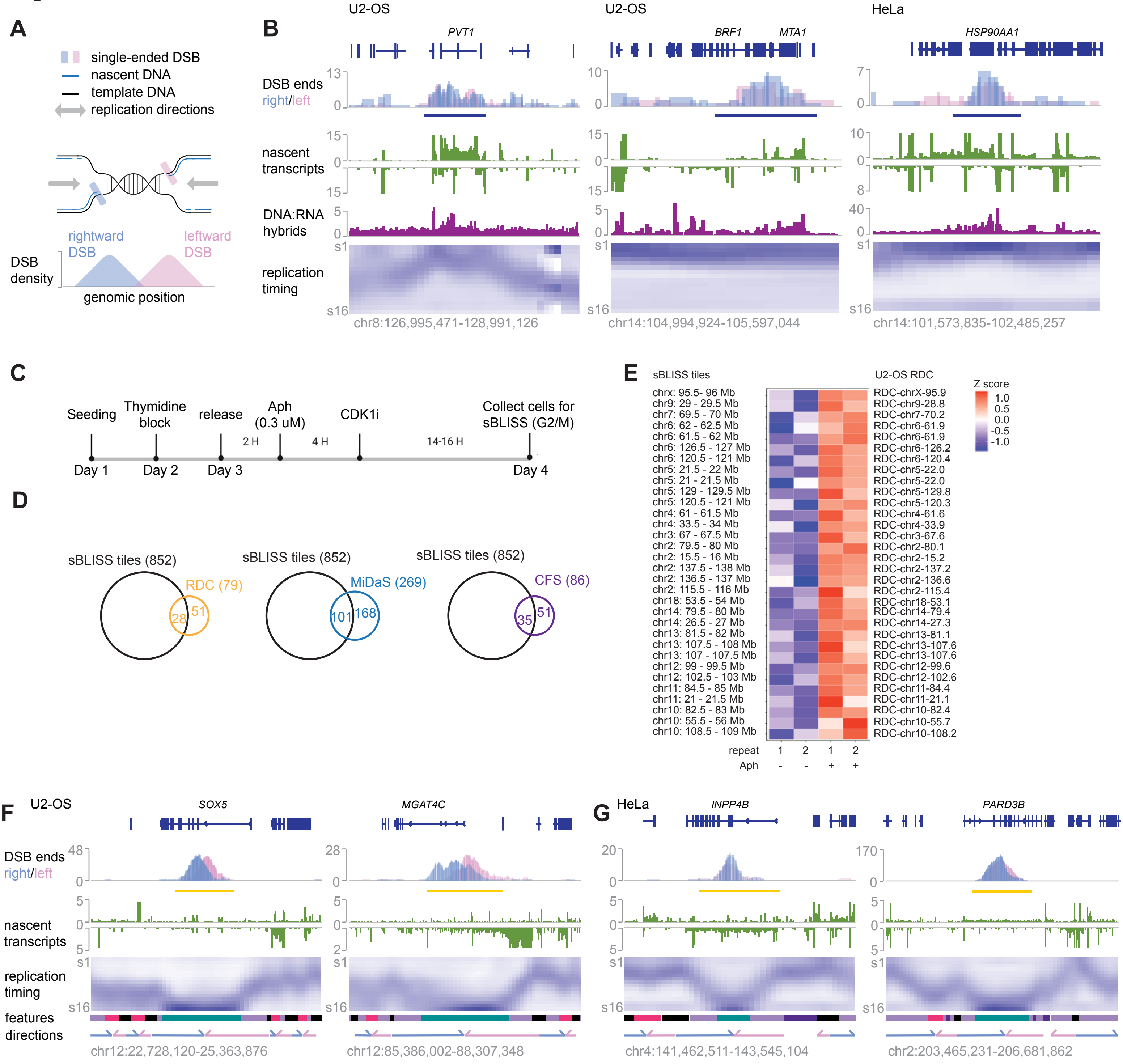

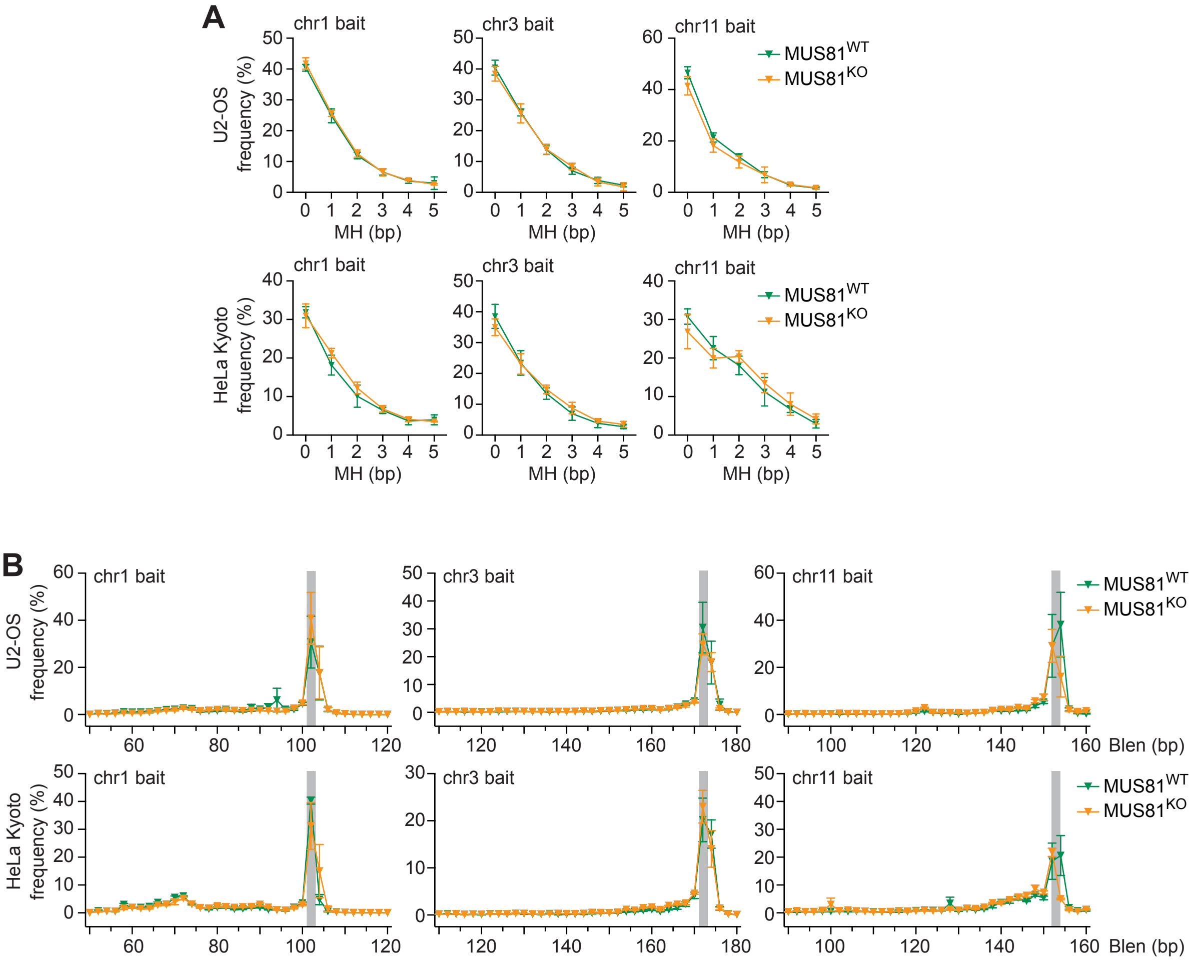

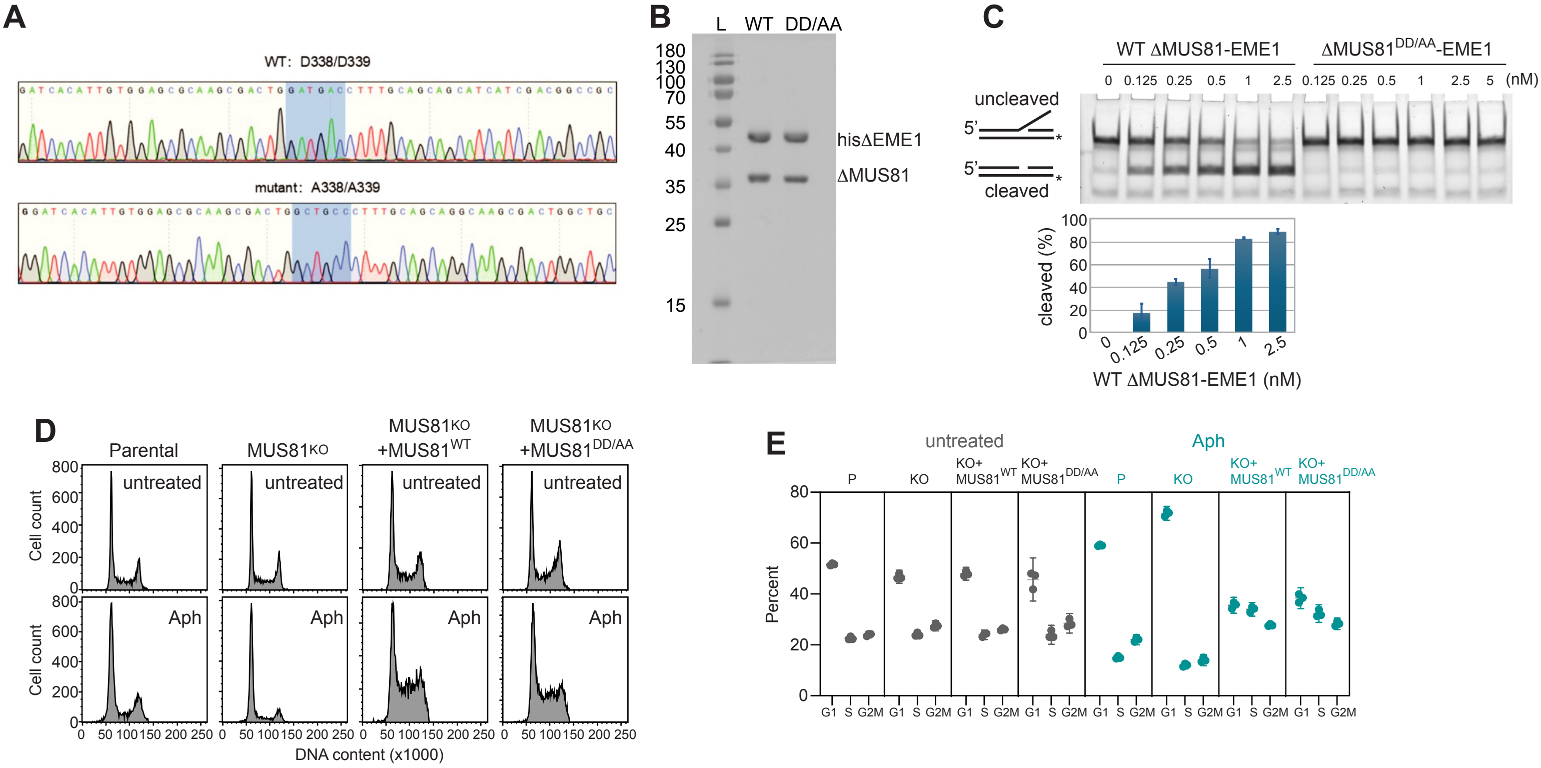

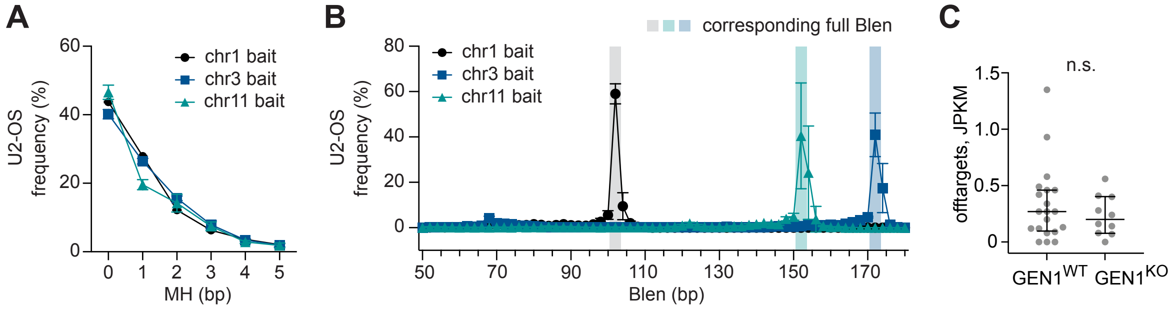

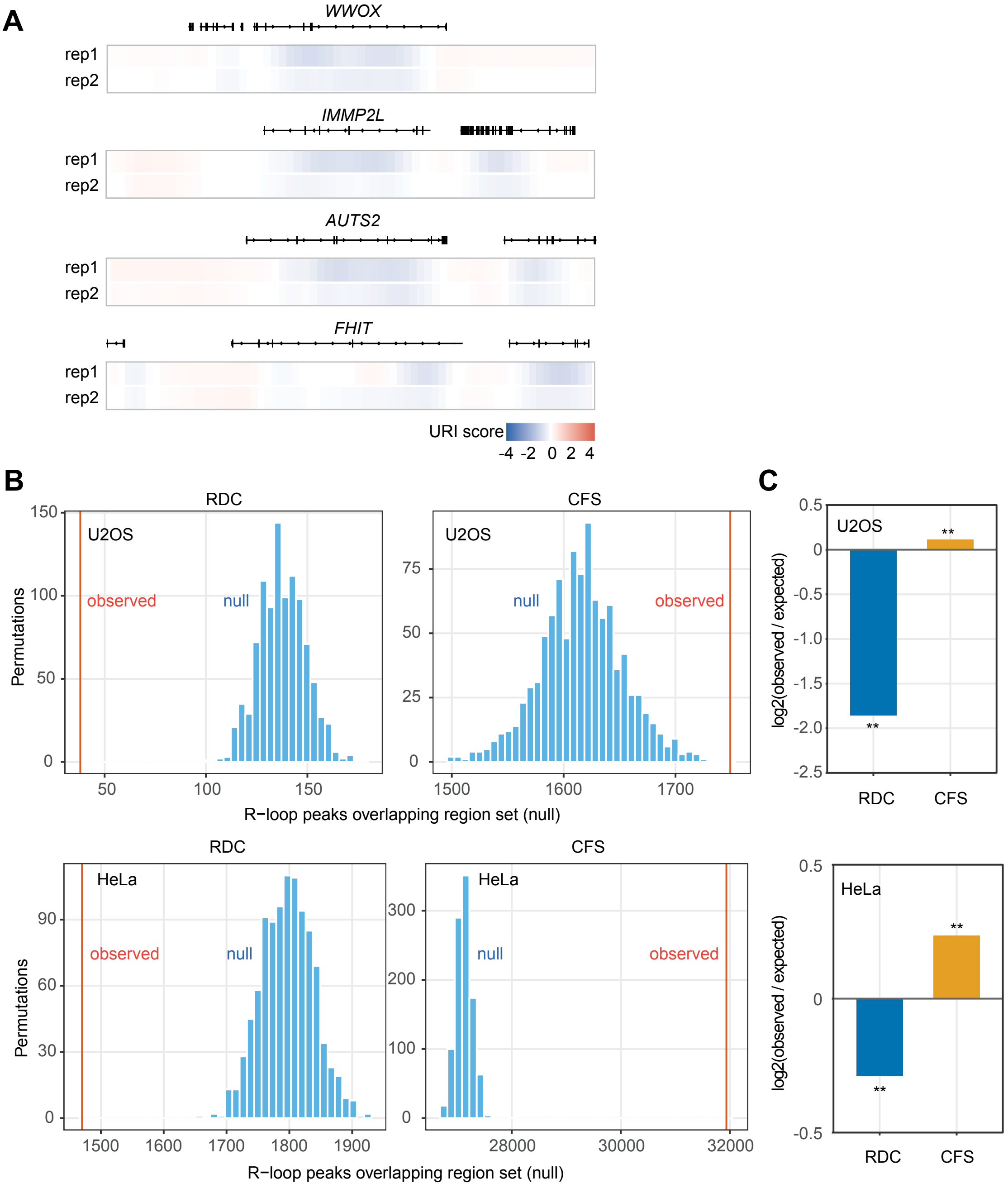
